# Mechanical activation of mitochondrial energy metabolism during cell differentiation

**DOI:** 10.1101/2022.03.23.485518

**Authors:** Zong-Heng Wang, Christian Combs, Wenjing Zhao, Jay Knutson, Mary Lilly, Hong Xu

## Abstract

In multicellular lives, differentiation of many types of stem and progenitor cells is often accompanied by a metabolic transition from glycolysis to mitochondrial oxidative phosphorylation. The mechanisms driving this metabolic transition in vivo are largely unknown. Here, we show that, during differentiation of the *Drosophila* female germline cyst, the surrounding somatic cells compress the cyst and increase the tension of cyst cells’ membranes. Transmembrane channel-like, an evolutionarily conserved ion channel involved in mechanosensation, maintains cytosolic Ca^2+^ levels in compressed differentiating cysts. Cytosolic Ca^2+^ induces transcriptional activation of oxidative phosphorylation through a CaMKI-Fray-JNK signaling relay. Our findings demonstrate a molecular link between cell mechanics and mitochondrial energy metabolism, with implications in other developmentally orchestrated metabolic transitions in mammals.

**One-Sentence Summary:** Mechanical forces from the surrounding tissue activate mitochondrial energy metabolism in differentiating cells in vivo.

## Main Text

Many types of stem cells and progenitor cells emphasize on glycolysis to preserve carbon sources for biosynthesis, while differentiated cells mainly rely on mitochondrial oxidative phosphorylation (OXPHOS), a more efficient way to produce ATP (*1, 2*). How mitochondrial energy metabolism is activated during cell differentiation in most developmental processes is unknown. Previous studies uncovered that OXPHOS is transcriptionally activated during differentiation of the *Drosophila* female germline cyst (Figure 1, A and B) (*3, 4*). In the *Drosophila* germarium, a cystoblast generated from asymmetric division of a germline stem cell undergoes 4 rounds of incomplete cytokinesis to form a cyst with 16 interconnected germ cells. Subsequently, the cyst is encased by somatic pre-follicle cells (pFCs), flattened into a one cell-thick disc, and starts to differentiate. The cyst that is consisted of an oocyte and fifteen nurse cells eventually becomes spherical in the budding egg chamber (Figure 1A) (*5-7*). The co-occurrence between OXPHOS activation and cyst shape remodeling intrigued us to test whether OXPHOS is induced in response to the flattening of differentiating cysts.

**Figure 1.**
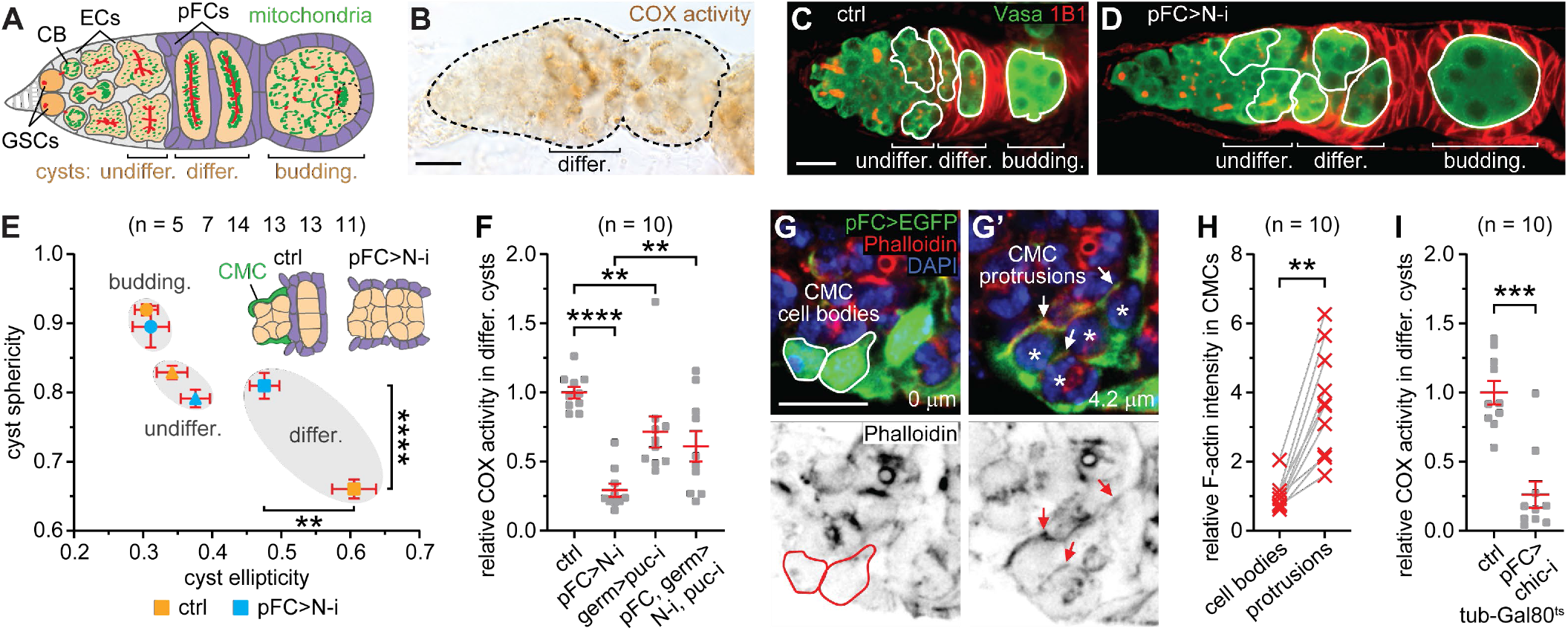
Cyst compression drives OXPHOS activation during differentiation. (**A**) Schematic of the germarium. GSCs, germline stem cells. CB, cystoblast. ECs, escort cells. pFCs, pre-follicle cells. (**B**) COX activity in a germarium. (**C** and **D**) Germaria stained for Vasa (germ cells) and 1B1 (pFCs’ cell membranes and fusomes). (**E**) Sphericity and ellipticity of developing cysts. CMC, cross-migrating pFC. (**F**) Quantification of COX activity in differentiating cysts. (**G** and **G’**) Two sections from a z-stack of CMCs in the germarium shown in fig. S2. Asterisks show germ cells in a cyst wrapped by CMCs. (**H**) Quantification of relative intensity of F-actin (Phalloidin) in 10 CMCs from 8 ctrl (control) germaria. (**I**) Quantification of COX activity in differentiating cysts. \*\**P* < 0.01, \*\*\**P* < 0.001, and \*\*\*\**P* < 0.0001 [Mann-Witney tests for (E), (F), and (I); Wilcoxon signed-rank test for (H)]. Scale bars, 10 µm.

Differentiating cysts might be flattened by the surrounding pFCs. Notch (N) signaling controls a few pFCs, called cross-migrating cells (CMCs), to migrate centripetally and position themselves between cysts (fig. S1A) (*8*). To test whether impairing pFCs’ migration could spare individual cysts from flattening, we inhibited N signaling in pFCs with an interfering RNA (RNAi) under the control of a pFC-Gal4 driver (109-30-Gal4) (*9*). The *N* RNAi disrupted pFCs’ migration (fig. S1B), phenocopying *N* mutation (*8*). Importantly, most differentiating cysts were no longer separated by pFCs and became rounded rather than flattened, both in a sagittal view and in a three-dimensional (3D) reconstruction, although they appeared differentiating normally (Fig. 1, C-E, fig. S1, B and C, and Movies S1 and S2). The shape of earlier- or later-stage cysts was not affected. These results indicate that differentiating cysts are compressed by the surrounding pFCs.

A JNK-induced insulin-Myc feedforward loop activates the transcription of OXPHOS genes in differentiating cysts (*3*). When N signaling was inhibited in pFCs, both JNK and OXPHOS activities were downregulated in differentiating cysts (Fig. 1F, and fig. S1, D and E), examined by a JNK activity reporter (*puc*-nLacZ; Puc is a target and a negative regulator of JNK signaling) and a cytochrome c oxidase (COX) activity histochemistry, respectively (*3, 10*). Importantly, *puc* RNAi in germ cells that can ectopically activate JNK signaling partially restored COX activity in the background of *N* RNAi in somatic pFCs (Fig. 1F), suggesting that active N signaling in pFCs is essential for JNK-mediated OXPHOS activation in germ cells. Besides promoting CMCs’ migration, N may also affect other aspects of interactions between pFCs and germ cells. We hence attempted to directly address whether pFCs’ cross-migration is necessary for OXPHOS activation in differentiating cysts. In line with other cell migration models (*11, 12*), when wrapping around the cyst, CMCs formed protrusions that accumulated more F-actin than the CMC cell bodies did (Fig. 1, G-H). F-actin is also enriched in the pFCs between differentiating cysts (fig. S2, A and A’), presumably providing rigidity for pFCs to compress cysts. Chickadee (Chic), the fly homolog of profilin that facilitates actin polymerization, is highly expressed in pFCs (*13, 14*). When Chic is reduced in pFCs, pFC’s positioning between cysts was impaired and differentiating cysts appeared partially relieved from compression, phenocopying the *N* RNAi (fig. S2B). Moreover, this genetic manipulation impaired OXPHOS activation in differentiating cysts (Fig. 1I). By contrast, depleting Egalitarian (Egl), a key regulator for cyst differentiation (*15*), barely affected OXPHOS in compressed cysts (fig. S3, A and B). Taken together, our results demonstrate that compression by the surrounding pFCs activates OXPHOS in differentiating cysts.

The fusome is a branched cytoskeleton-rich structure extending into all cyst cells through the actin-rich intercellular bridges, ring canals (*16-18*). During cyst compression, the fusome may resist the sliding and twisting between cyst cells and pull the membranes at the cyst cell interfaces, thereby locally stretching their membranes (Fig. 2A). To test this idea, we performed fluorescence lifetime imaging microscopy on living ovaries labelled with Flipper-TR, a cell membrane tension reporter (*19*). Increased membrane tension enlarges the spaces between phospholipid tails and allows Flipper-TR to adopt a trans-conformation that shortens its lifetime (*19*). As cysts differentiated, Flipper-TR lifetime became shorter at the cyst cells’ interfaces but remained the same at the cyst’s borders. Relieving differentiating cysts from compression with *N* RNAi in pFCs abolished the shortening of Flipper-TR lifetime at the cyst cells’ interfaces (Fig. 2, B and C, and fig. S4, A and B). Because cysts do not grow during differentiation (fig. S1F), cyst compression is likely the driving force behind the expansion of cyst cells’ interfaces.

**Figure 2.**
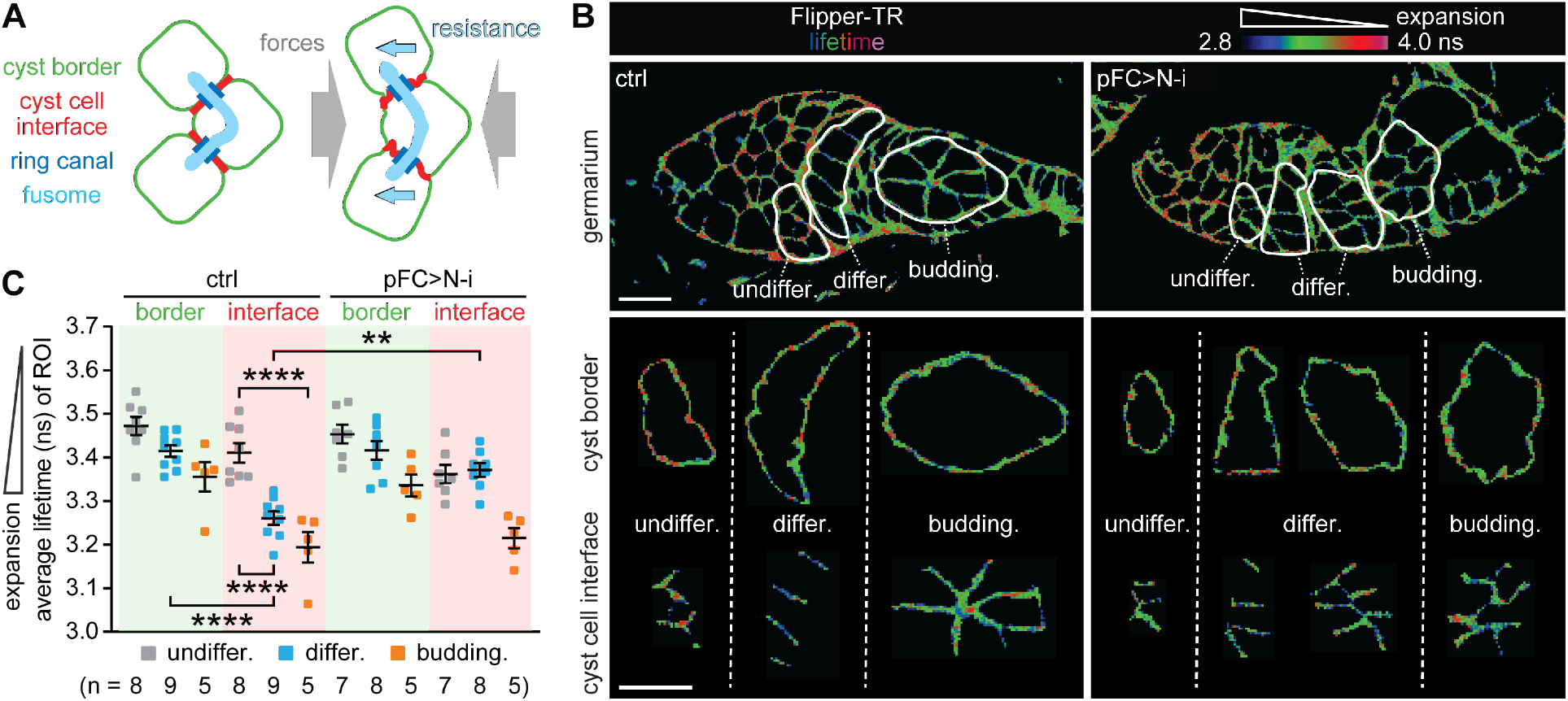
Cell membranes are expanded in compressed differentiating cysts. (**A**) Cyst cells twist and slide relatively to each other during cyst compression. Resistance from the fusome may stretch membranes at the cyst cell interfaces. (**B**) Images of Flipper-TR lifetime at cell membranes in germaria. (**C**) Quantification of average Flipper-TR lifetimes in regions of interest (ROI). \*\**P* < 0.01 and \*\*\*\**P* < 0.0001 (Mann-Witney test). Scale bars, 10 µm.

Mechanoresponsive molecules on the cell membrane transduce mechanical stimuli to modulate various cellular processes (*20-22*). To test whether any mechanoresponsive molecules is involved in OXPHOS activation in differentiating cysts, we performed a candidate germline RNAi screen of 40 genes annotated as “responsive to mechanical stimuli” for the phenotype of reduced COX activity (Table S1). RNAi against transmembrane channel-like (Tmc) strongly impaired COX activity in differentiating cysts, as well as JNK activation and Myc stabilization (Fig. 3A, and fig. S5). In the background of germline *Tmc* RNAi or *tmc*^*1*^ mutation, germline *puc* RNAi restored most COX activity (Fig. 3B and fig. S5B). Tmc is an evolutionarily conserved ion channel mediating mechanosensation in response to cell membrane expansion (*23, 24*), suggesting that the expansion of cyst cell’s membranes is sensed by Tmc and subsequently activates OXPHOS through JNK.

**Figure 3:**
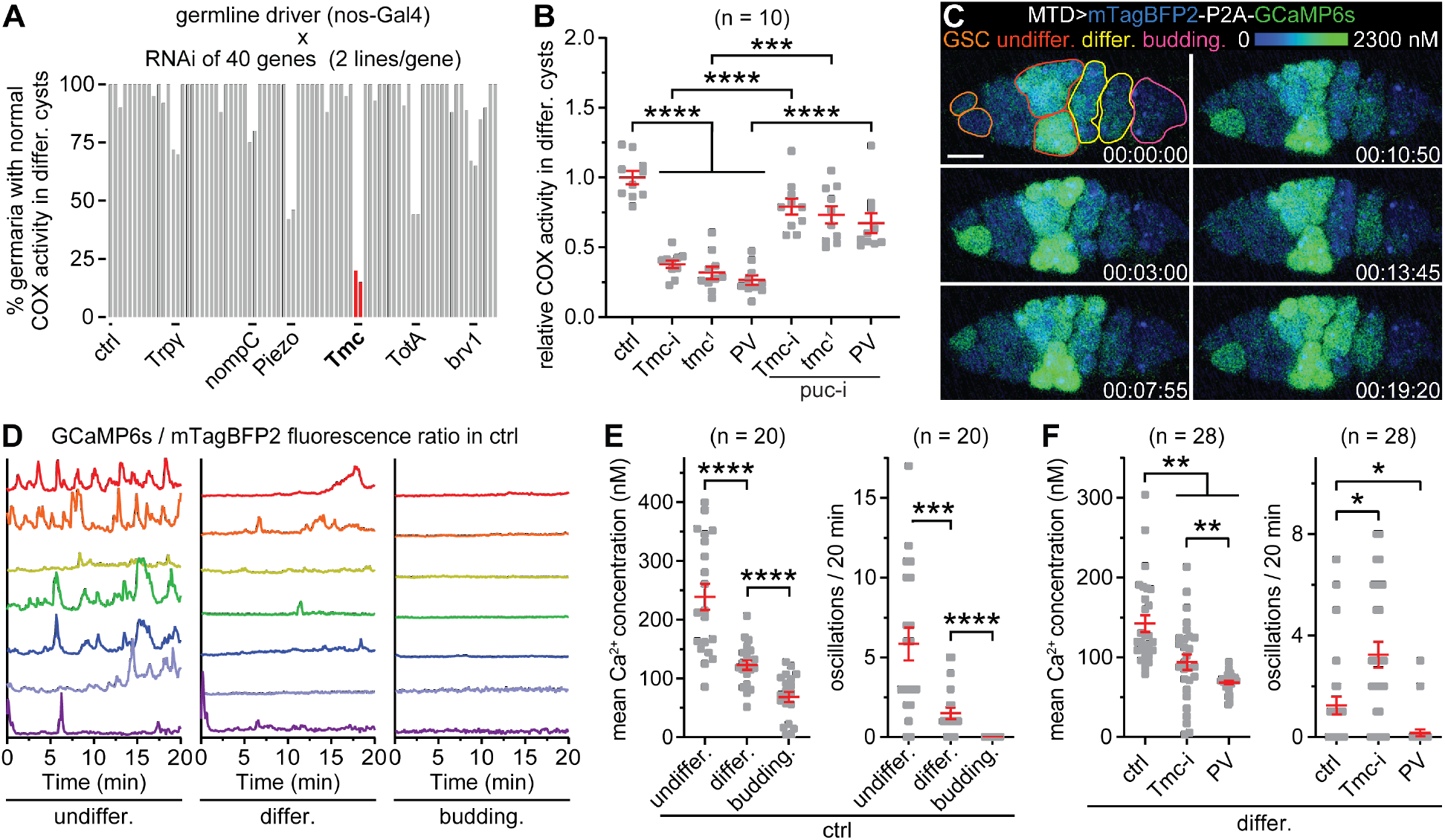
Tmc mediates OXPHOS activation by maintaining cytosolic Ca^2+^ concentrations. **(A)** A germline RNAi screen for mechanoresponsive genes. (**B**) Quantification of COX activity in differentiating cysts. (**C**) Live recording showing Ca^2+^ levels and oscillations in germ cells at different stages (time stamp in h:min:s). The range for intracellular Ca^2+^ concentration is shown. (**D**) Representative traces of GCaMP6s/mTagBFP2 fluorescence ratio in individual ctrl germ cells at different developmental stages from 5 germaria. (**E** to **F**) Mean Ca^2+^ concentrations and Ca^2+^ oscillation frequency in individual germ cells from 5 germaria for (E) and 7 germaria for (F). \**P* < 0.05, \*\**P* < 0.01, \*\*\**P* < 0.001, and \*\*\*\**P* < 0.0001 [Kruskal-Wallis tests for the four groups in (B) and the three groups in (F, left); Mann-Witney tests between the two groups in (B), (E), (F), and (G)]. Scale bar, 10 µm.

The opening of Tmc channel allows the influx of Ca^2+^ into the cytosol (*25, 26*), prompting us to examine the cytosolic Ca^2+^ profile along cyst development. We generated a UASz transgene that expresses a fusion protein consisted of mTagBFP2 and GCaMP6s linked by a self-cleaving P2A peptide (fig. S6A). GCaMP6s is a highly sensitive fluorescence Ca^2+^ reporter (*27*), while mTagBFP2 serves as the internal control to normalize the expression levels of GCaMP6s (fig. S6B). We calibrated this ratiometric Ca^2+^ reporter in perfused S2 cells using a series of Ca^2+^ buffers (fig. S6C). Simultaneous live imaging for both fluorescent proteins in germaria revealed highly variable Ca^2+^ concentrations and distinct patterns of oscillations in germ cells at different developmental stages (Fig. 3C). In the control germarium, undifferentiated cysts had higher cytosolic Ca^2+^ levels (∼239 nM) and Ca^2+^ oscillation frequency (5.8 times per 20 min), compared to differentiating cysts (∼123 nM and 1.5 times per 20 min) and budding egg chambers (69 nM and 0 time per 20 min) (Fig. 3, D and E, and Movie S3). *Tmc* RNAi had no obvious influence on either cytosolic Ca^2+^ concentrations or oscillations in undifferentiated cysts, presumably due to high Ca^2+^ concentrations at this stage (fig. S7, A and B, and Movies S4). In differentiating cysts, *Tmc* depletion lowered Ca^2+^ concentrations but increased Ca^2+^ oscillation frequency (Fig. 3F, fig. S7 A and C, and Movies S4). To further understand whether either or both of these two abnormalities may account for the OXPHOS defect in *Tmc* depleted differentiating cysts, we ectopically expressed a high-affinity Ca^2+^ binding Parvalbumin (PV) to sequestrate free intracellular Ca^2+^ (*28*). Compared to *Tmc* RNAi, PV expression led to a greater reduction in Ca^2+^ concentrations and diminished Ca^2+^ oscillations in differentiating cysts, while it similarly impaired JNK and COX activities (Fig. 3, B and F, fig. S7, fig. S8, A and B, and Movies S5). Also, the COX defect could be rescued by *puc* RNAi in both *Tmc* RNAi and PV expressing cysts (Fig. 3B). Therefore, the role of Tmc seems to maintain cytosolic Ca^2+^ concentrations that leads to OXPHOS activation through JNK, while the frequency of Ca^2+^ oscillations is not critical. As *Tmc* transcripts were uniformly expressed along cyst differentiation (fig. S9, A and B), other mechanisms must contribute to the higher basal Ca^2+^ levels in undifferentiated cysts. Although mitochondrial Ca^2+^ regulates cell metabolism (*29*), knockdown of the mitochondrial uniporter (MCU) had little impact on COX activity (fig. S8, C and D), further substantiating that cytosolic Ca^2+^ plays a critical role in mechanical activation of OXPHOS during cyst differentiation.

Despite high levels of cytosolic Ca^2+^, JNK and OXPHOS are not activated in undifferentiated cysts. We therefore hypothesized that the signaling molecules, other than JNK (fig. S9, A and B), that function downstream of Ca^2+^ to activate OXPHOS in differentiating cysts might be absent in undifferentiated cysts. To this end, we assessed the potential involvement of a number of genes related to Ca^2+^ signaling and JNK pathway in OXPHOS activation in differentiating cysts (Table 2). Germline RNAi of Calcium/calmodulin-dependent protein kinase I (CaMKI) or Frayed (Fray), a conserved kinase regulating osmolarity (*30*), abolished JNK and COX activities in differentiating cysts. Moreover, COX activity in either *CaMKI* or *Fray* RNAi was partially restored by *puc* RNAi (Fig. 4A and fig. S10, A to D). Importantly, endogenous expression of GFP-tagged CaMKI was low in early cysts but markedly increased in differentiating cysts and thereafter, while the levels of endogenously expressed Fray-GFP remained nearly uniform (Fig. 4B and fig. S11A). Thus, the developmental patterns of CaMKI likely confine cytosolic Ca^2+^-induced JNK activation to differentiating cysts.

**Figure 4:**
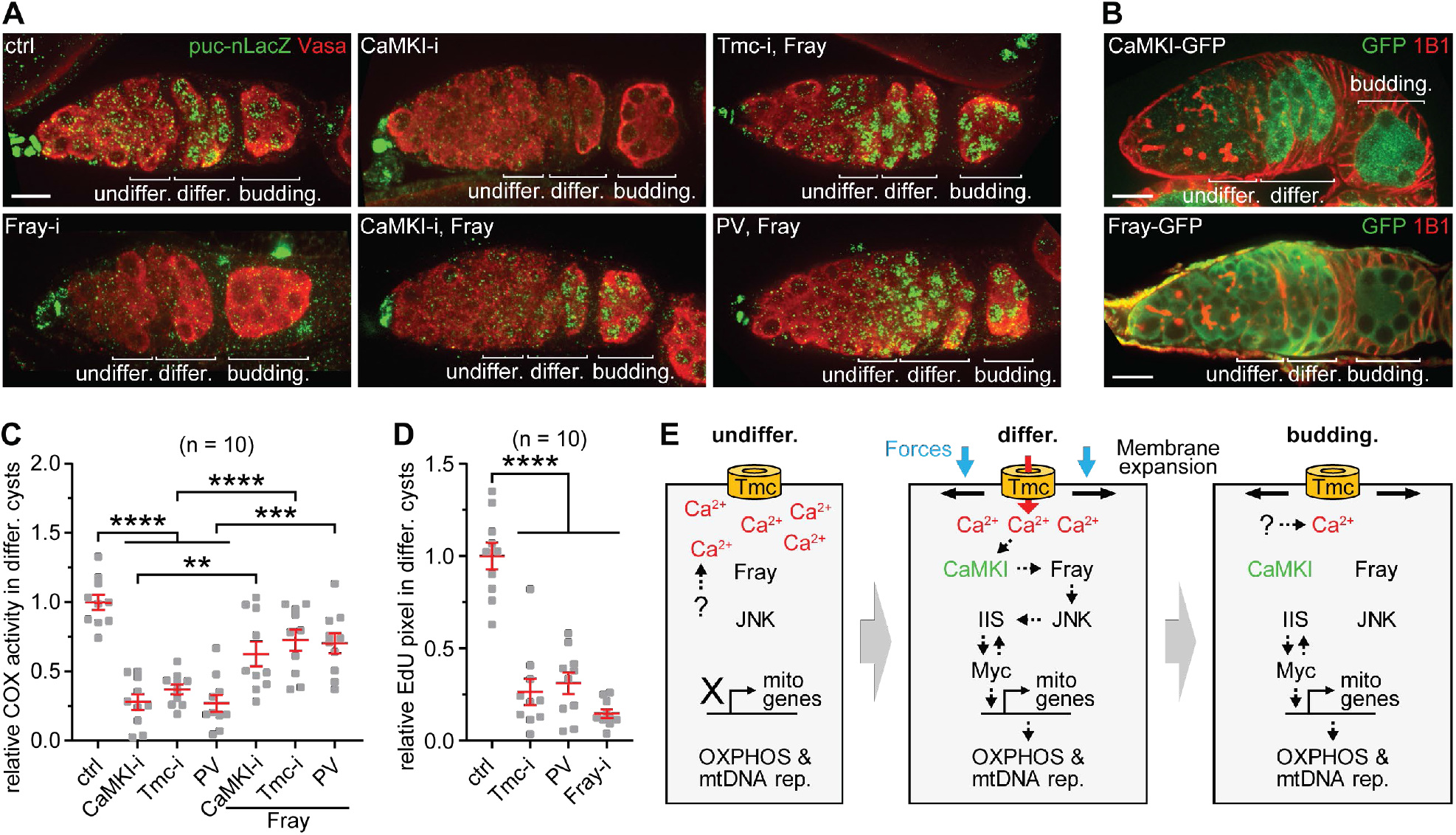
Cytosolic Ca^2+^ mediates mechanical activation of OXPHOS in differentiating cells through CaMKI and Fray. (**A**) Visualization of JNK activity in germaria. (**B**) Expression of endogenous CaMKI-GFP and Fray-GFP in germaria. (**C**) Quantification of COX activity in differentiating cysts. (**D**) Quantification of EdU incorporated into mtDNA in differentiating cysts. (**E**) Schematic of the mechanotransduction pathway that activates OXPHOS during cyst differentiation. \*\**P* < 0.01, \*\*\**P* < 0.001, and \*\*\*\**P* < 0.0001 [Kruskal-Wallis tests for the four groups in (C) and (D); Mann-Witney tests between the two groups in (C)]. Scale bars, 10 µm.

To determine the epistasis between Fray and CaMKI, we generated a UASz transgene (UASz-Fray-3HA) expressing a wild-type, active Fray kinase (*31*). Overexpression of Fray enhanced both JNK and COX activities in differentiating cysts with *Tmc* or *CaMKI* RNAi, or overexpressing PV, and ectopically induced JNK activity in the budding egg chamber (Fig. 4, A and C, and fig. S11B). Therefore, Fray mediates Ca^2+^ and CaMKI in JNK and OXPHOS activation during cyst differentiation.

JNK-induced OXPHOS supports mtDNA replication during *Drosophila* oogenesis and maintains fertility (*3*). Consistent with our proposal that Tmc maintains cytosolic Ca^2+^ to activate JNK and OXPHOS, *Tmc* RNAi, *Fray* RNAi or PV overexpression markedly impaired mtDNA replication in the germline and caused reduced egg hatching rate (Fig. 4D and fig. S11, C and D).

Unicellular eukaryotes modulate energy metabolism directly in response to nutrient availability in the environment, while stem cells and their differentiated progeny in multicellular organisms often reside in the same environment with a controlled supply of nutrients (*32*). How OXPHOS activation is coupled with cell differentiation remains an enigma. In this study, we uncover that mechanical forces from the surrounding tissue function as developmental cues to activate mitochondrial energy metabolism during differentiation of the *Drosophila* female germline cyst (Fig. 4E). During neuronal differentiation in the mammalian cerebral cortex, differentiating neurons migrate towards the cortical plate by squeezing through layers of cells while increasing their OXPHOS activity (*33, 34*). Moreover, maturing cardiomyocytes (CMs) feature increased Ca^2+^-dependent contractility and enhanced OXPHOS activity (*35, 36*). CM contraction is modulated by stretch-activated ion channels and several CaMK genes are upregulated during CM maturation (*37, 38*). Our work establishes a molecular link between cell mechanics and mitochondrial energy metabolism, which may represent an evolutionarily conserved mechanism in animal development.

## Acknowledgements

We thank F. Chanut for comments and editing on the manuscript; the Bloomington Drosophila Stock Center and the Vienna Drosophila Resource Center for providing fly lines; the Developmental Studies of Hybridoma Bank for various antibodies; Bestgene Inc. for Drosophila transgenesis service.

## Fundings

This work is supported by the National Heart Lung and Blood Institute Intramural Research Program and the Eunice Kennedy Shriver National Institute of Child Health and Human Development Intramural Research Program.

## Author contributions

Conceptualization: Z.-H.W., H.X.; Investigation: Z.-H.W., C.C.; Methodology: Z.-H.W., C.C., W.Z., J.K.; Funding acquisition: J.K., M.L., H.X.; Project administration: Z.-H.W., H.X.; Supervision: Z.-H.W., M.L., H.X.; Visualization: Z.-H.W., C.C., W.Z., H.X.; Writing - original draft: Z.-H.W., C.C., H.X.; Writing - review and editing: Z.-H.W., M.L., H.X.

## Competing interests

The authors declare no competing interests.

## Data and materials availability

All data are available in the main text or the supplementary materials. Materials are available upon request.

## Supplementary Materials

### Materials and Methods

#### Fly stocks

Flies were maintained on standard BDSC cornmeal medium at 25°C. RNAi lines for candidate screen are listed in Tables S1 and S2. *Luciferase*-RNAi (ctrl, BL31603), *109-30*-Gal4 (soma-Gal4, BL7023), *nos*-Gal4 (germline, BL25751), *nos*-Gal4 (germline, BL32563), UAS-2xEGFP (BL6874), *tub*-Gal80^ts^ (BL7018), *MTD*-Gal4 (germline, BL31777), *puc*-nLacZ (*puc*-lacZ^A251.1F3^, BL11173), NRE-EGFP (BL30728), *N*-RNAi (BL27988), *chic*-RNAi (BL34523), *tmc*^*1*^ (BL66556) (*1*), *puc*-RNAi (BL36085), *egl*-RNAi (BL43550), *MCU*-RNAi (BL67857), *CaMKI*-RNAi (BL35362), *CaMKI*-RNAi #2 (BL41900), *fray*-RNAi (BL42569), *fray*-RNAi #2 (BL55878), and Myc-GFP (BL81274) were obtained from the Bloomington Drosophila Stock Center. CaMKI-GFP (v318349) and Fray-GFP (v318460) were from the Vienna Drosophila Resource Center. Vasa-GFP (#109171) was purchased from the Kyoto *Drosophila* Genomics and Genetics Resources.

#### Female fly hatching rate

Virgin female flies with different genotypes were crossed with young (3-4 days) *w*^*1118*^ male flies in batches overnight. More than 180 embryos were collected and separated into 4 groups on grape juice agar plates with a small amount of yeast paste at the center. Plates were placed in a 25°C incubator for 2 days. The hatching rate was calculated as the number of hatched larvae divided by the number of embryos, averaged across the 4 groups.

#### Transgenic flies

To generate pUASz-mTagBFP2-P2A-GCaMP6s, GCaMP6s was amplified from pGP-CMV-GCaMP6s (#40753, addgene) and mTagBFP2 from pCAG-mTagBFP2 (*2, 3*). GCaMP6s and mTagBFP2, as well as a P2A fragments (GCCACCAACTTCTCCCTGCTGAAGCAGGCCGGCGACGTGGAGGAGAACCCCGGCCCC), were subcloned into XhoI-cut UASZ-1.0 (*4*) with the In-Fusion Cloning kit (Takara Bio Inc.). The PV DNA fragment and the coding region of Fray were amplified from pCMV-PV-GFP (#17301, addgene) and RE53265 (BDGP cDNA library), respectively. PV and Fray, as well as a C-terminal 3xHA tag, were subcloned into XhoI-cut UASZ-1.0 with the In-Fusion Cloning kit. Primers were listed in Table S3. Transgenic flies carrying these constructs were generated by Bestgene Inc.

#### Immunofluorescence staining and EdU labeling

Immunofluorescence staining and EdU labeling were performed as previously described (*5*). For immunofluorescence staining, antibodies/dye used were as follows: rabbit anti-Vasa (1:1000, Santa Cruz Biotechnology), rabbit anti-GFP (1:1,000, NB600-308, Novus Biologicals); mouse anti-Orb (1:800, 6H4, Developmental Studies Hybridoma Bank); mouse anti-Hts (1:800, 1B1, Developmental Studies Hybridoma Bank); mouse anti-β-galactosidase (1:1200, Z378A, Promega); Alexa Fluor 568-Phalloidin (1:500, A12380, Invitrogen); Alexa Fluor 568 donkey anti-rabbit IgG (1:600, A10042, Invitrogen), Alexa Fluor 568 goat anti-mouse IgG (1:600, A11004, Invitrogen), Alexa Fluor 488 goat anti-rabbit IgG (1:600, A11034, Invitrogen), and Alexa Fluor 488 goat anti-mouse IgG (1:600, R37120, Invitrogen).

The Click-iT EdU labeling kit (C10637, Invitrogen) was utilized to label EdU according to manufacturer’s instructions. For EdU incorporation assay, ovaries were dissected in Schneider medium supplemented with 10% FBS (A3840002, Gibco) and pretreated with 7 µM aphidicolin (A0781,Sigma-Aldrich) in the medium for 3 h at room temperature. Then, the ovaries were incubated with 10 µM EdU and 7 µM aphidicolin in the medium with 10% FBS for 2 h. After three washes in the medium, ovaries were fixed in 4% PFA in PBS for 20 min, washed twice in PBS with 3% BSA for 5 min, and permeabilized in PBS with 0.5% Triton X-100 for 30 min. Subsequently, the ovaries were washed twice in PBS and co-stained with anti-1B1.

Samples were mounted with VECTASHIELD mounting medium with DAPI (H-1500, Vector Laboratories). Confocal images of germaria were collected on a Perkin Elmer Ultraview system (Zeiss Plan-apochromat 63×/1.4 oil lens, Volocity software, Hamamatsu Digital Camera C10600 ORCA-R2, Immersol immersion oil 518F) with a 0.3 μm step size. Images for morphological analysis of female germline cysts were acquired with an Instant Sim (iSIM) Super-Resolution Microscope (Olympus UPlanSApo 60×/1.30 Sil lens, Metamorph acquisition software, ORCA-Flash4.0 V2 Digital CMOS camera C11440, Silicone Immersion Oil SIL300CS-30SC) with a 0.2 μm step size. All confocal images were processed with NIH ImageJ (National Institutes of Health).

#### Morphological analysis of female germline cysts in 3D

Z-stack images were opened with Imaris (version 9.7, Oxford Instruments). Individual 16-cell cysts were identified based on DAPI staining. In germaria with somatic *N* RNAi, fused cysts were observed in some budding egg chambers. We only analyzed the shape of budding egg chamber cysts with 16 cells. In Imaris, 3D surface view of individual germline cyst was generated with the ‘surfaces’ tool. ‘Draw’ was used to manually trace the same cyst in each confocal step. After generating 3D surfaces with ‘create surface’, the values of cyst sphericity, ellipticity (oblate), and size were obtained in ‘detailed statistics’. Images of 3D surfaces were generated with ‘snapshot’ and 3D movies were created with ‘animation’.

#### Single molecule fluorescence in situ hybridization (smFISH)

Fluorescently (Quasar® 570) labeled Stellaris FISH probes against *Tmc*, and *JNK* mRNA were synthesized by Biosearch Technologies. smFISH on *Drosophila* ovaries were performed according to a published protocol (*6*). Briefly, dissected Vasa-GFP expressing ovaries were fixed in 4% PFA in PBS for 20 min. After 5×5 in washes with PBST (PBS and 0.2% Triton X-100), ovaries were permeabilized with 3 µg/ml proteinase K in PBS on ice for 1 h. Permeabilization was stopped by incubating ovaries in PBS with 20 mg/ml glycine and was followed by a post-fixation in PBST with 4% PFA for 20 min. After 5×2 min washes with PBST, ovaries were incubated in prehybridization solution (2× SSC and 10% formamide) at room temperature for 10 min. Prehybridization solution was removed, and 60 µl of hybridization solution (6 µl deionized formamide, 3 µg heparin, 1 µl salmon sperm DNA, 80 ng probe mix, 6 µg dextran sulfate, 120 µg BSA, and 5 µl of 20× SSC) was added. Ovaries were protected from light and incubated at 37°C overnight. Ovaries were then washed for 2×15 min with prewarmed prehybridization solution in at 37°C, followed by 2×5 min washes with PBS at room temperature. Z-stack images with smFISH, Vasa-GFP, and nuclear (DAPI) channels were collected with the Perkin Elmer Ultraview system (Zeiss Plan-apochromat 63×/1.4 oil lens, Volocity software, Hamamatsu Digital Camera C10600 ORCA-R2, Immersol immersion oil 518F) and processed with NIH ImageJ. The sequence of fluorescently labeled short DNA probes targeting *Tmc* and *JNK* mRNA are listed in Table S4.

#### Calcium imaging

Three-day-old females were collected. Their ovaries were dissected in Schneider medium (Sigma-Aldrich) and immerged in hydrogel as adapted from (*7*). Briefly, dissected ovaries were transferred into a droplet of medium on a 22×22 mm coverslip that were previously coated with 3-(trimethoxysilyl) propyl methacrylate (440159, Sigma Aldrich). Medium was then replaced by 15 µl of 10% PEG-DA hydrogel solution (GS700, Advanced BioMatrix, Inc.) with 0.1% Irgacure 2959 (photo initiator, 410896, Sigma-Aldrich). A coverslip treated with deperlent was placed above the hydrogel solution. The coverslip/coverslip sandwich was illuminated by a UV light source for 30 s at 312 nm for gelation. The upper coverslip was removed. The bottom coverslip with the hydrogel was placed into a Chamlide chamber (CM-S22-1, Quorum Technologies) filled with Schneider medium. Time-lapse imaging (every 5 s for 20 min) of the middle section (z-axis) of the germarium was performed on a Leica SP8 confocal microscope (HC PL APO CS2 63×/1.40 N.A. objective lens) using the Las X software (version 3.5.7). Fluorescence of GCaMP6s (excitation: 488 nm, emission: 500-550 nm, and gain: 150%) and mTagBFP2 (excitation: 405 nm, emission at 440-480 nm, and gain: 0%) were recorded simultaneously as videos at 8-bit depth.

The video flies were opened with NIH ImageJ. ‘Freehand selections’ was used to select germ cell areas. Time-lapse fluorescence intensities for both channels in individual germ cells were obtained with ‘plot z-axis profile’. Time-lapse background intensities of each channel calculated from a 10×10 µm^2^ square outside the germarium were subtracted from the time-lapse fluorescence intensities. Time-lapse ratios of GCaMP6s/mTagBFP2 intensities were imported into Origin 2021 (OriginLab Corporation). In Origin 2021, Ca^2+^ oscillation in each germ cell during the 20-min recording were counted manually based on the individual peaks of GCaMP6s/mTagBFP2 intensity ratio. In Excel, average Ca^2+^ level for each germ cell was calculated by averaging the fluorescence ratio of the 20-min imaging. The amplitude of oscillation spikes was calculated by subtracting the average ratio of two timepoints flanking the peak from the altitude of the peak obtained in Origin 2021. The baseline Ca^2+^ level between oscillations was calculated by averaging the ratio without the values of peaks.

For Ca^2+^ calibrations, the mTagBFP2-P2A-GCaMP6s fragment was firstly subcloned into the pIB/V5-His vector (V802001, Invitrogen) with the In-Fusion Cloning kit. 2×10^6^ S2 cells (*Drosophila* Genomics Resource Center, S2-DRSC, from Nasser Rusan’s lab) were seeded in a 60 mm dish and transfected with 2 µg the pIB-mTagBFP2-P2A-GCaMP6s-V5-His construct using Effectene Transfection Reagent (Qiagen). After 2 days, cells were spread on concanavalin A-coated chambered coverglasses (155411, Thermo Scientific) and permeabilized by 150 μM digitonin (D141, Sigma-Aldrich) in the zero free calcium buffer (30 mM MOPS, pH 7.2, 100 mM KCl, 10 mM EGTA; see below) for 10 min. A series of buffers with 11 different free calcium concentrations were obtained by mixing the zero free calcium buffer (30 mM MOPS, pH 7.2, 100 mM KCl, 10 mM EGTA) and the 39 μM free calcium buffer (30 mM MOPS, pH 7.2, 100 mM KCl, 10 mM CaEGTA) in various ratios (Calcium Calibration Buffer Kit #1, Life Technologies) and added to different wells of chambered coverglasses. Confocal images of transfected cells (n = 13∼29 cells for each calcium concentration) were acquired by a Leica SP8 confocal microscope (HC PL APO CS2 63x/1.40 N.A. objective lens) using the Las X software (version 3.5.7) with the same settings for Ca^2+^ imaging on germaria. In Prism 9 (GraphPad), background removed ratios of GCaMP6s/mTagBFP2 intensities were plotted to generate a sigmoidal standard curve that was then utilized to fit germ cell data.

#### Membrane tension measurements

Cell membrane tension was measured with Flipper-TR fluorescent tension probe (SC020, Cytoskeleton, Inc.) (*8*). Dissected ovaries were incubated with 2 µM Flipper-TR in Schneider medium for 30 min. Single z-plane FLIM images of the germarium regions were acquired using a Leica SP8 Falcon FLIM confocal microscope, a HC PL APO CS2 63×/1.40 N.A. objective lens, and Leica Las X (version 3.5.7). Point scanning excitation at a speed of 400 Hz was performed at 488 nm using a pulsed white-light laser operating at 80 MHz with emission collected over a bandwidth of 550-650 nm onto a hybrid single molecule detector (HyD SMD) at 16-bit digitization with a pinhole set to 1 A.U. Image size was set to 512×512 pixels^2^ with pixel sizes of 300 nm. Fluorescence from germaria as frame accumulated over 70 images to build up an adequate number of photons per pixel for further analysis. Time-correlated single photon counting histograms were collected with 136 channels in a 13 ns time window (97 ps per channel). Before lifetime fitting, pixel binning of 2 was performed to provide peak counts of at least 600 photons/pixel for the dual exponential fits, followed by a manual thresholding to omit background pixels. Fluorescence decay data from full images was fitted to a dual exponential tail fit model in the Las X software. Images of the longest lifetime were exported for further analysis. A custom program written in IDL (Interactive Data Language, L3Harris Geospatial) was used to obtain average lifetime values in regions of interest (ROIs) from the lifetime pixel-map images generated by the Las X software.

#### COX activity histochemistry of fly ovaries

Histochemical activity staining and quantification for cytochrome c oxidase (complex IV) were performed according to the procedure from our previous study (*5*). Four to eight pairs of ovaries from 2∼3 day-old flies were dissected in PBS and ovarioles were separated by using a dissection needle. Ovaries were incubated with COX staining solution [50 mM phosphate solution (pH 7.4), 4 mM 3,3’-diaminobenzidine, 2 µg/ml catalase, 200 μM cytochrome c, 4 mM antimycin A, 84 mM malonate, and 60 µM rotenone] for 30 min at room temperature. For negative control, ovaries (*Luciferase* RNAi) were treated with 2 mM KCN in COX staining solution for 30 min at room temperature. Negative control was performed with each batch of COX activity staining. All reactions were followed by 2×5 min washes with phosphate solution and 4% paraformaldehyde fixation for 15 mins. After 2×5 min washes in phosphate solution, ovaries were immersed in 80% glycerol in phosphate solution. Brightfield images of germaria were collected by a Zeiss Axio Observer Z1 microscope (C-Apochromat 40×/1.1 W Corr objective lens for ovaries).

Relative ETC activities in differentiating cysts were quantified with ImageJ. From the opened images, areas of differentiating cysts were isolated with ‘freehand selections’ and followed by ‘clear outside’. The color of the cysts was converted into gray and inverted to obtain images with black background. The COX staining area was selected with ‘color threshold’. Mean intensity of the selected COX staining area was measured. Mean intensity of non-selected area was considered as background and subtracted from the mean intensity of selected area. From each batch of activity staining, relative COX activity of ctrl differentiating cysts was considered as ‘1’, while the intensity from the negative control was considered as ‘0’ activity.

#### Quantification and statistical analysis

Sample size was not predetermined by statistical methods. The experiments were not randomized. Investigators were not blinded. Prism 9 (GraphPad) was used to plot data and perform statistical analyses. Error bars in all charts represent standard errors. Mann-Whitney test (two-tailed) was used to determine the mean differences between two unpaired groups, and Wilcoxon signed-rank test (two-tailed) for the differences between two paired groups. Kruskal-Wallis test, followed by Dunn’s multiple comparisons test, was performed to compare three and more groups. Differences were considered statistically significant when *P* < 0.05.

## Supplementary Figures and Legends

**Fig. S1.**
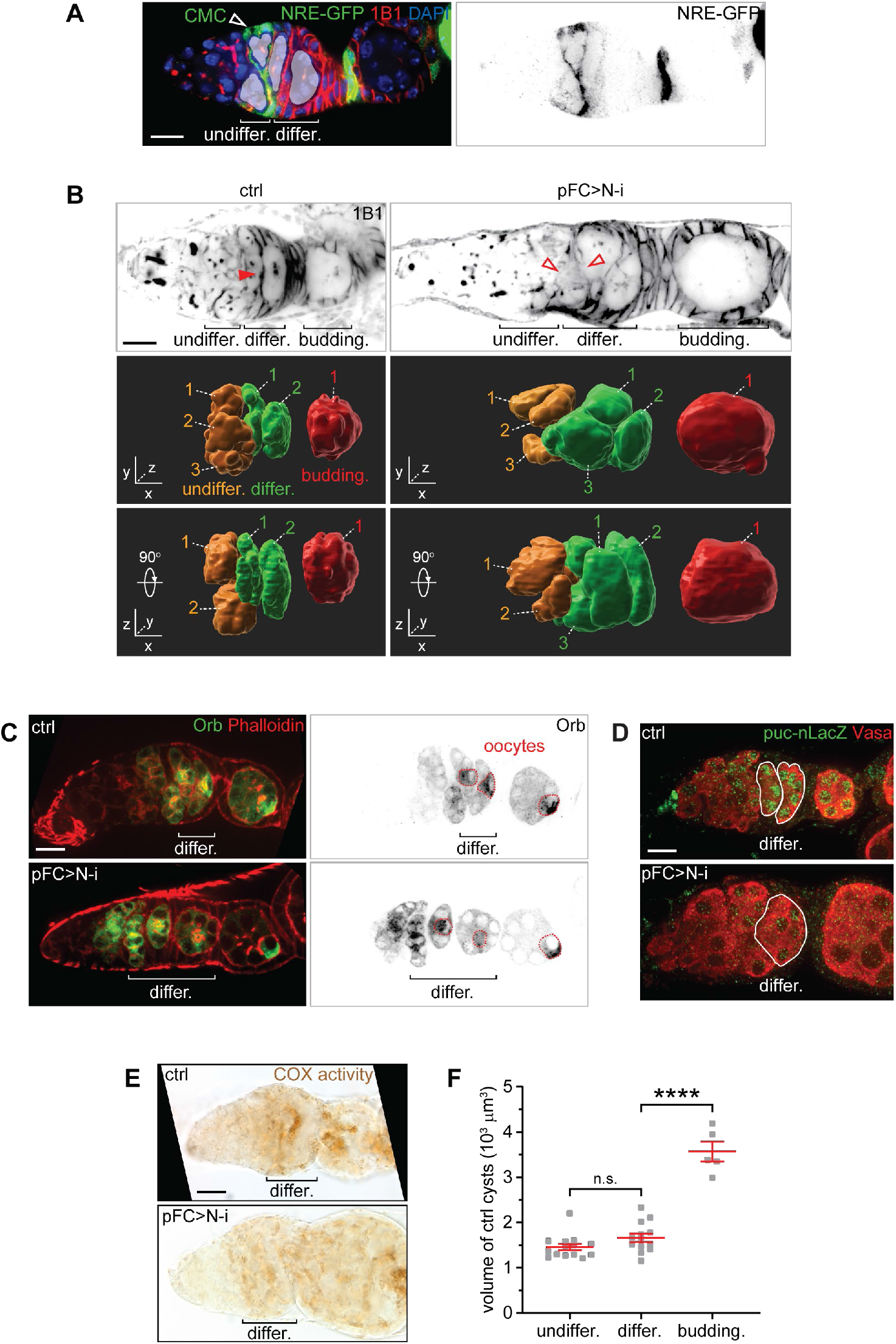
pFCs’ wrapping is required for compression and OXPHOS induction, but not differentiation, of the female germline cysts. (**A**) Representative image of Notch signaling activity (NRE-GFP) in the germarium. CMC, cross-migrating pFC. (**B**) 1B1 staining and three-dimensional reconstructions of the 16-cell cysts from Fig. 1, C and D. Red Arrow head points to a pFC inserted between germline cysts. Hollow arrow heads indicate that pFCs are absent between cysts. (**C**) Cyst differentiation visualized by Orb (oocyte marker) immunostaining. (**D**) Visualization of JNK activity by *puc*-nLacZ in germaria. (**E**) Representative images of COX activity histochemistry in differentiating cysts. (**F**) Cyst sizes in ctrl germaria. n = 14 (undifferentiated cysts), n = 13 (differentiating cysts), and n = 5 (budding egg chamber cysts) from 5 ctrl germaria. \*\*\*\**P* < 0.0001 and n.s., not significant (Mann-Witney test). Scale bars, 10 µm.

**Fig. S2.**
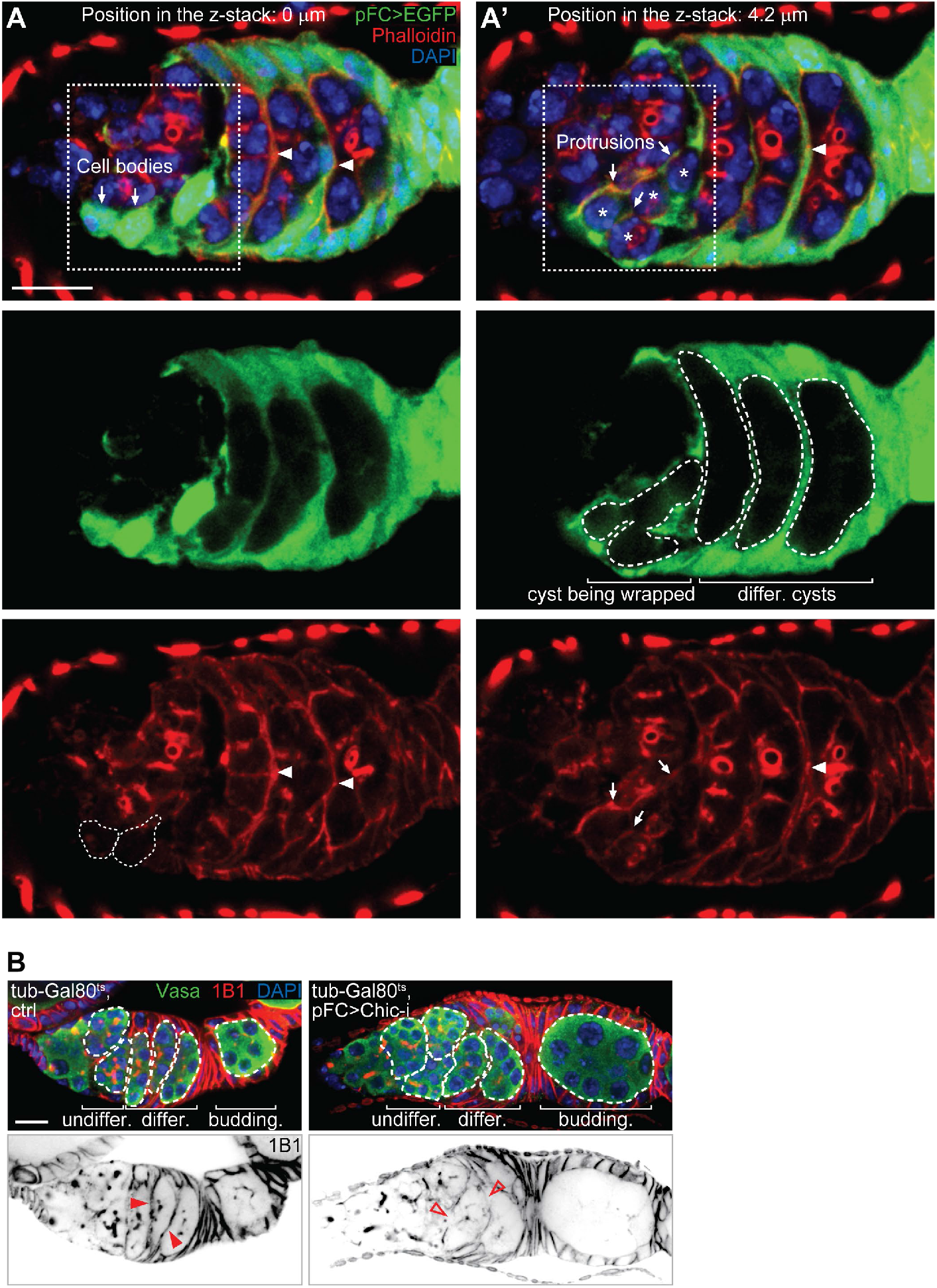
F-actin in CMCs and the pFCs between differentiating cysts. (**A** and **A’**) Representative images of a germarium expressing EGFP in pFCs. Arrows indicate CMC cell bodies or protrusions. Asterisks show the nuclei of cells in a cyst wrapped by CMCs. Arrowheads point to the pFCs between differentiating cysts. Germline cysts are outlined in the EGFP image, CMC cell bodies are outlined in the phalloidin image. (**B**) Germaria stained for Vasa, 1B1, and DAPI. Red arrow head points to pFCs inserted between germline cysts. Hollow arrow heads indicate that pFCs are absent between cysts. Scale bars, 10 µm.

**Fig. S3.**
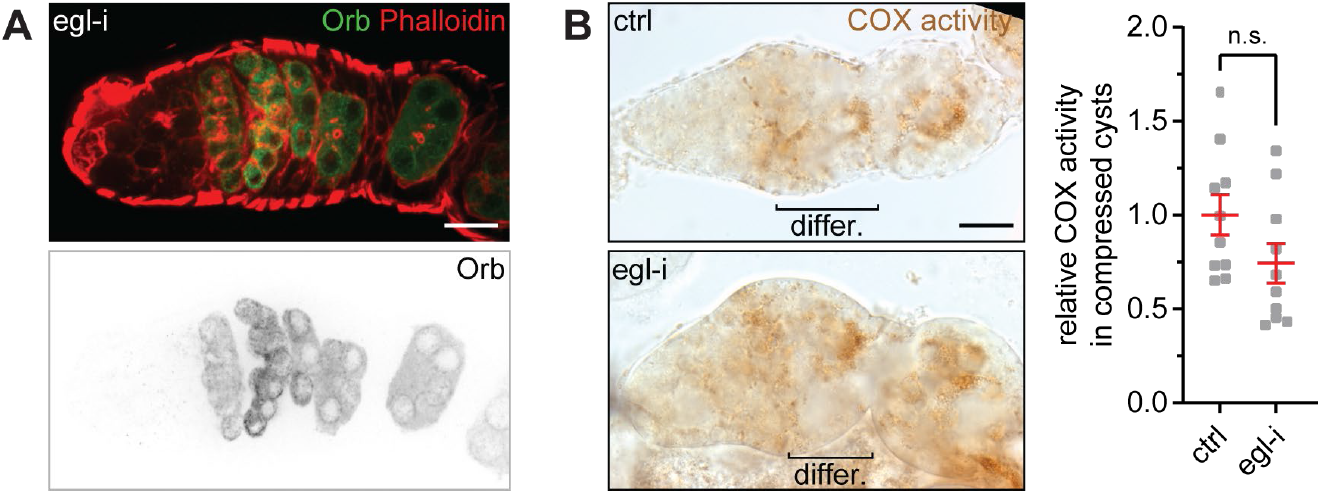
OXPHOS induction is independent of cyst differentiation. (**A**) Cyst differentiation is disrupted by *egl* RNAi. (**B**) COX activity in compressed cysts. n = 10 germaria for each genotype. n.s., not significant (Mann-Witney test). Scale bars, 10 µm.

**Fig. S4.**
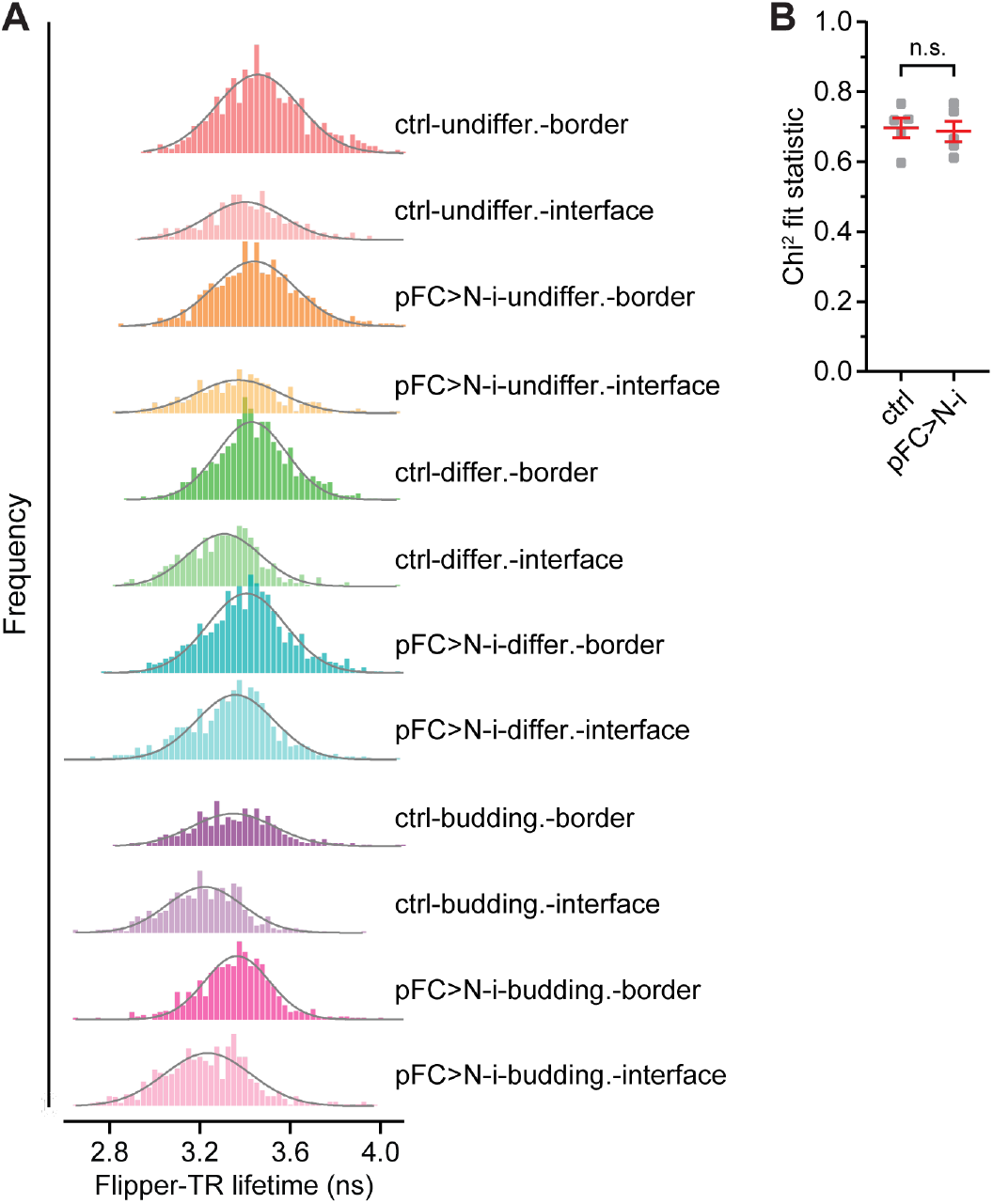
Cell membranes are expanded in compressed differentiating cysts. (**A**) Frequency distribution of Flipper-TR fluorescence lifetimes with Gaussian fits in ROIs. n = 8, 8, 7, 7, 9, 9, 8, 8, 5, 5, 5, 5 cysts, respectively. (**B**) Chi^2^ measurements for goodness of fits for the single images of germaria stained with Flipper-TR membrane tension probe. n.s., not significant (Mann-Witney test).

**Fig. S5.**
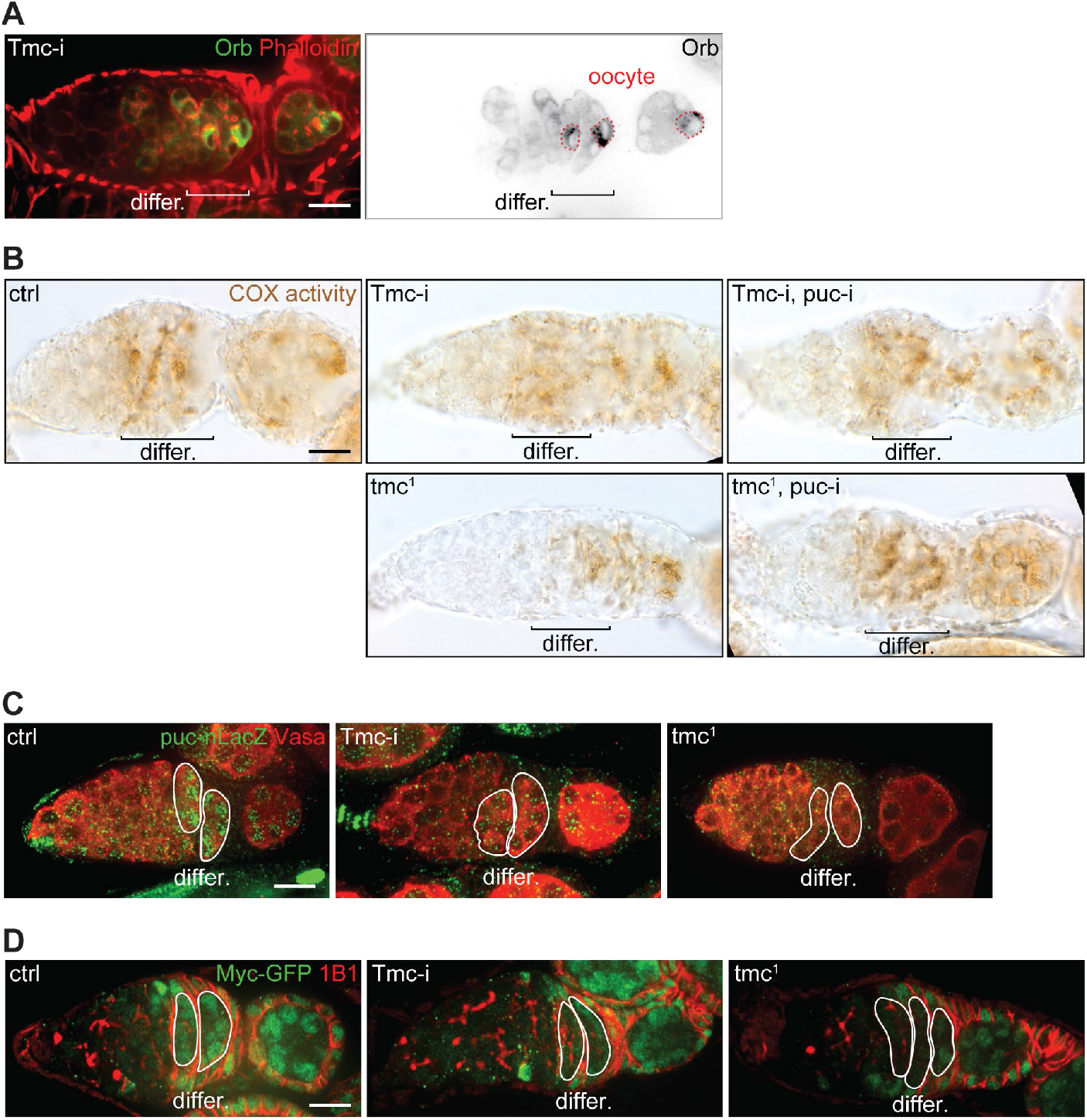
OXPHOS induction in differentiating cysts requires Tmc. (**A**) Cyst differentiation is normal in *Tmc* RNAi germarium. (**B**) COX activity histochemistry in germaria. (**C**) Representative images of JNK activity in germaria visualized by *puc*-nLacZ. (**D**) Representative images of endogenous Myc-GFP expression in germaria. Scale bars, 10 µm.

**Fig. S6.**
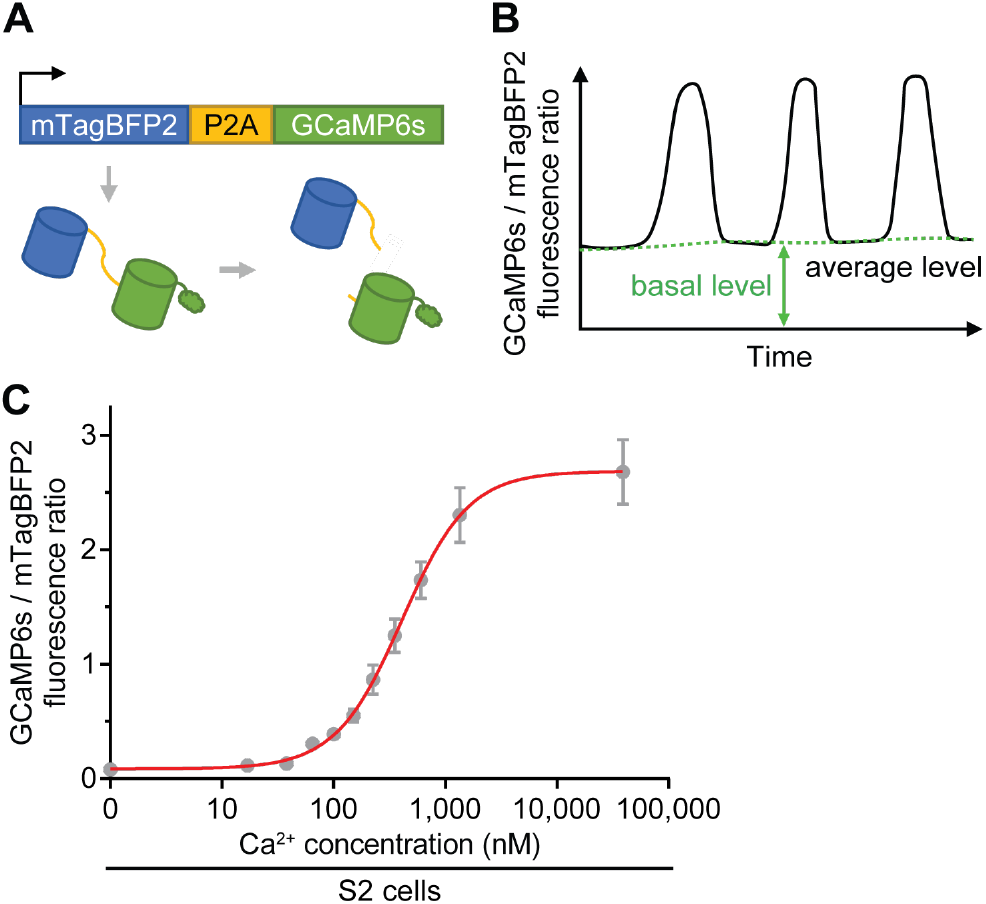
The UASz-mTagBFP2-P2A-GCaMP6s fusion protein and its calibration in S2 cells. (**A**) Schematic of UASz-mTagBFP2-P2A-GCaMP6s. (**B**) Ca^2+^ oscillation parameters. (**C**) Ca^2+^ calibrations of the mTagBFP2-P2A-GCaMP6s fusion protein in S2 cells. The standard curve is shown in red.

**Fig. S7.**
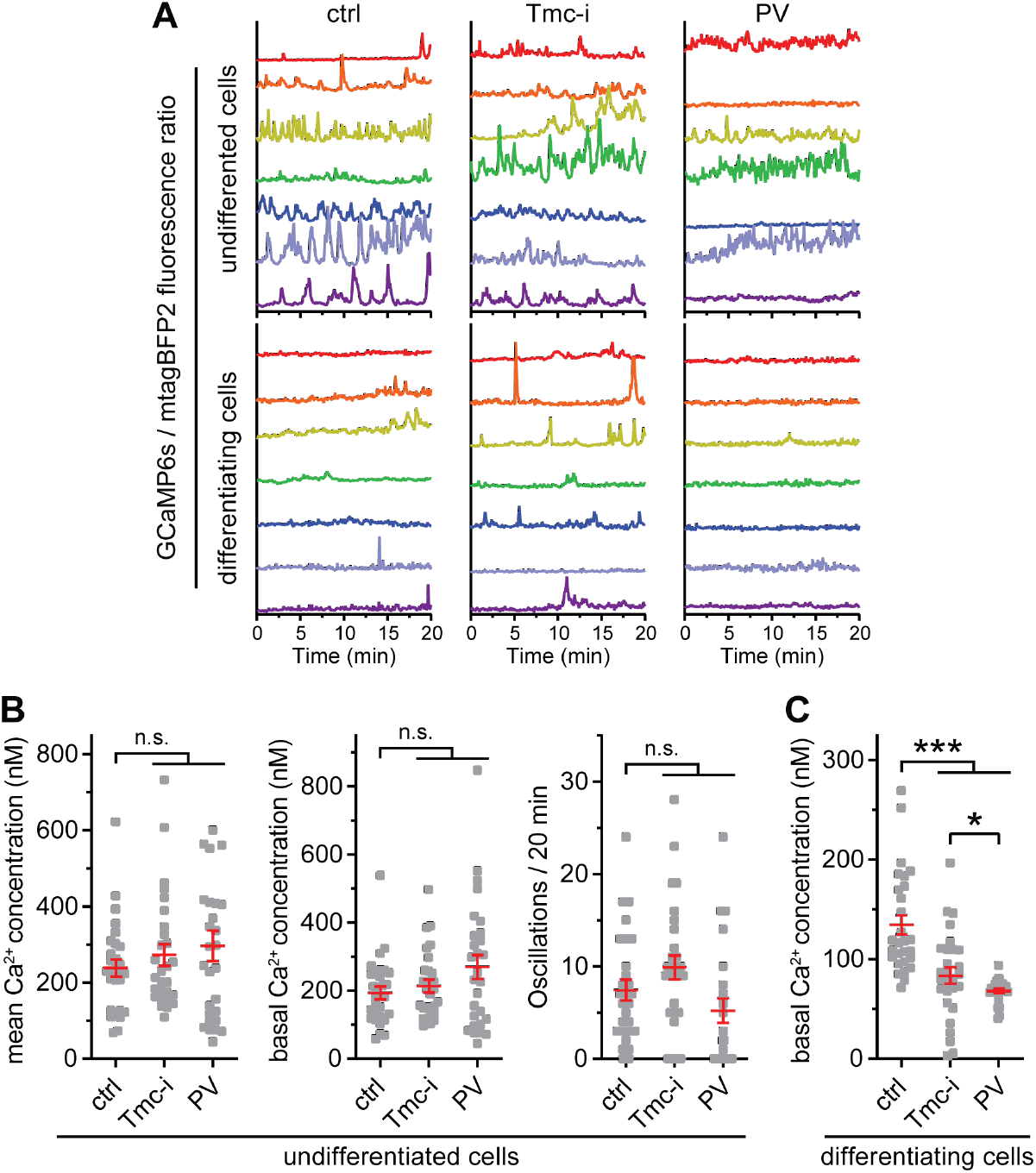
Cytosolic Ca^2+^ in differentiating but not undifferentiated cysts can be manipulated by *Tmc* knockdown or PV expression. (**A**) Representative traces of GCaMP6s/mTagBFP2 ratio in individual germ cells at different developmental stages. n = 7 germaria for each genotype. (**B** and **C**) Mean and basal Ca^2+^ concentrations, and Ca^2+^ oscillation frequency in individual germ cells. n = 28 cells from 7 germaria for each genotype. \**P* < 0.05, \*\*\**P* < 0.001, and n.s., not significant [Kruskal-Wallis tests for the three groups in (**B**) and (**C**); Mann-Witney tests between the two groups in (**C**)]. Scale bars, 10 µm.

**Fig. S8.**
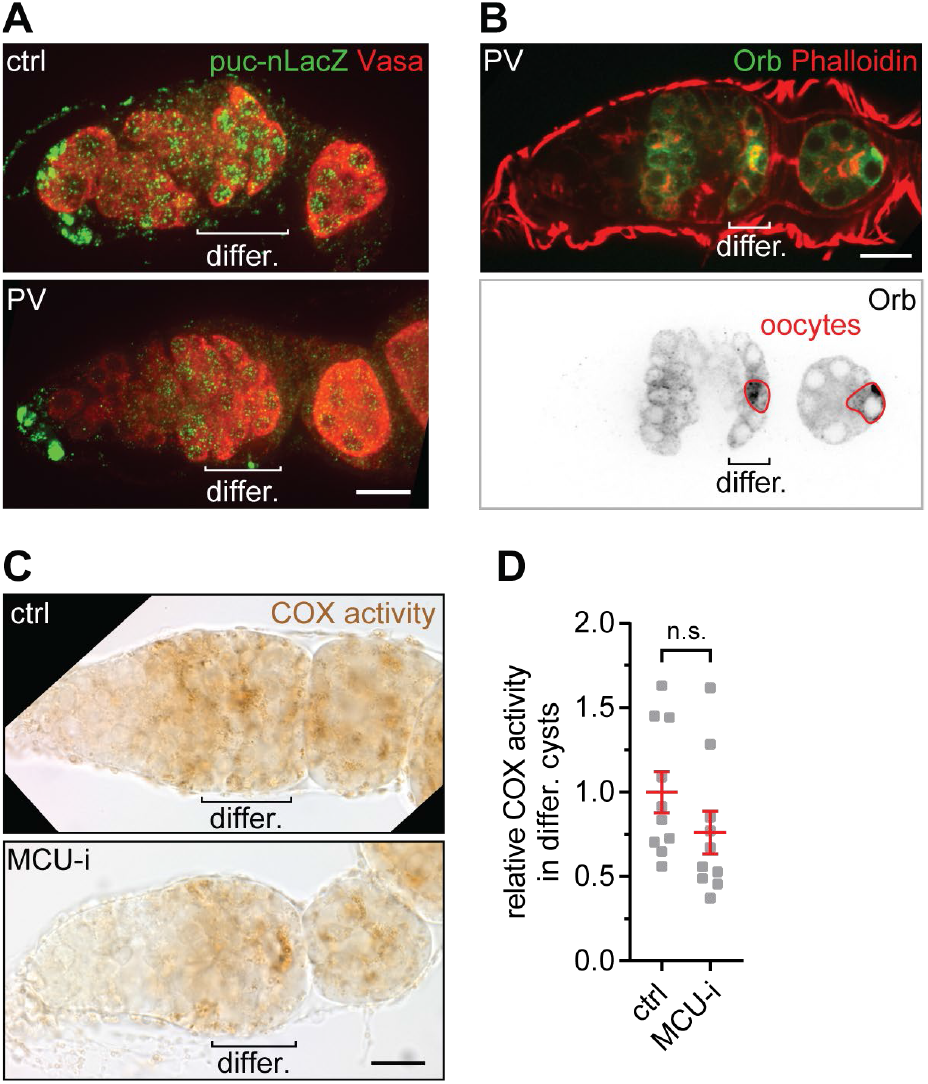
Cytosolic but not mitochondrial Ca^2+^ is essential for JNK and OXPHOS induction in differentiating cysts. (**A**) JNK activity is diminished in germ cells with expression of PV. (**B**) Cyst differentiation is normal when PV is expressed in the germline. (**C**) COX activity histochemistry in germaria. (**D**) Quantification of COX activity in differentiating cysts. n = 10 germaria for each genotype. n.s., not significant (Mann-Witney test). Scale bars, 10 µm.

**Fig. S9.**
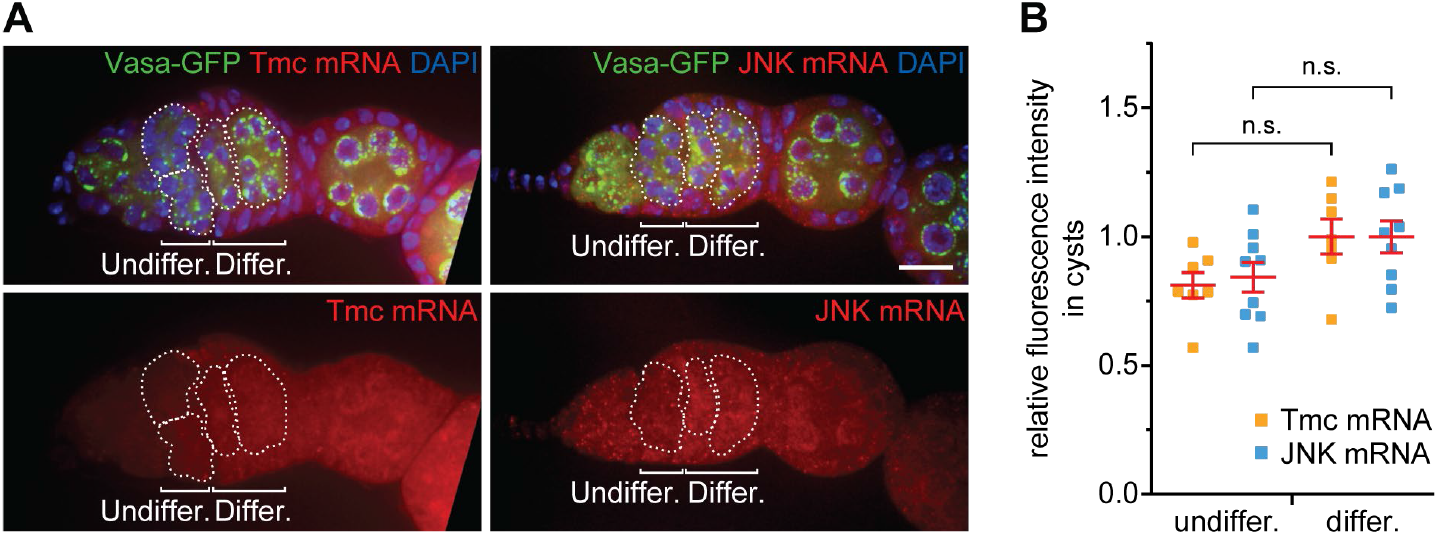
Expression of *Tmc* and *JNK* mRNA in germaria. (**A**) Visualization of *Tmc* and *JNK* mRNA in germaria by smFISH. (**B**) Quantification of *Tmc* (n = 7 germaria) and *JNK* mRNA (n = 9 germaria) in cysts. n.s., not significant (Mann-Witney tests). Scale bars, 10 µm.

**Fig. S10.**
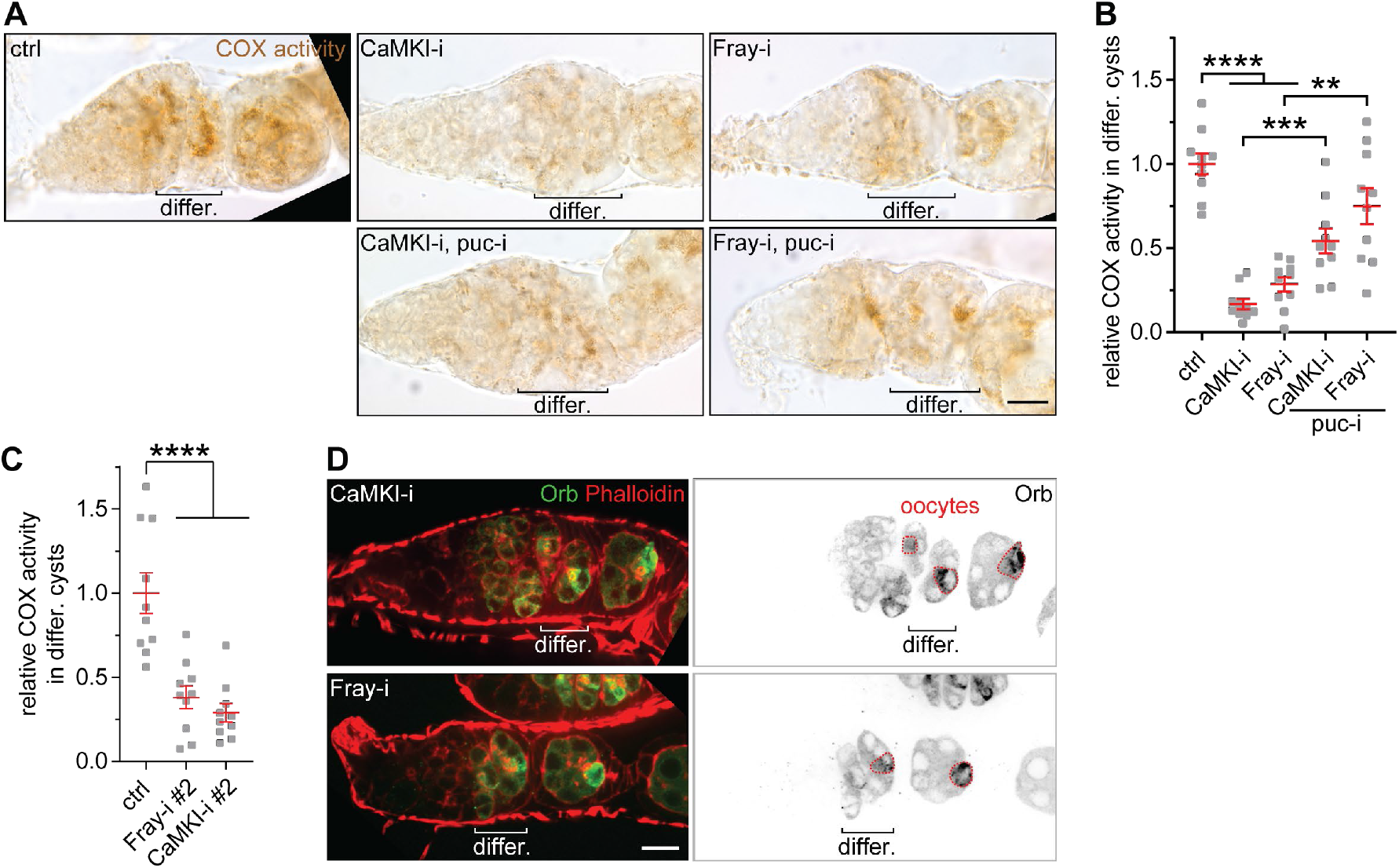
CaMKI and Fray are required for OXPHOS induction in differentiating cysts. (**A**) Representative images of COX activity histochemistry in germaria. (**B** and **C**) Quantification of COX activity in differentiating cysts. n = 10 germaria for each genotype. (**D**) Cyst differentiation is normal in *CaMKI* RNAi and *Fray* RNAi germaria. \*\**P* < 0.01, \*\*\**P* < 0.001, and \*\*\*\**P* < 0.0001 [Kruskal-Wallis tests for (**B**) and (**C**); Mann-Witney tests for (**B**)]. Scale bars, 10 µm.

**Fig. S11.**
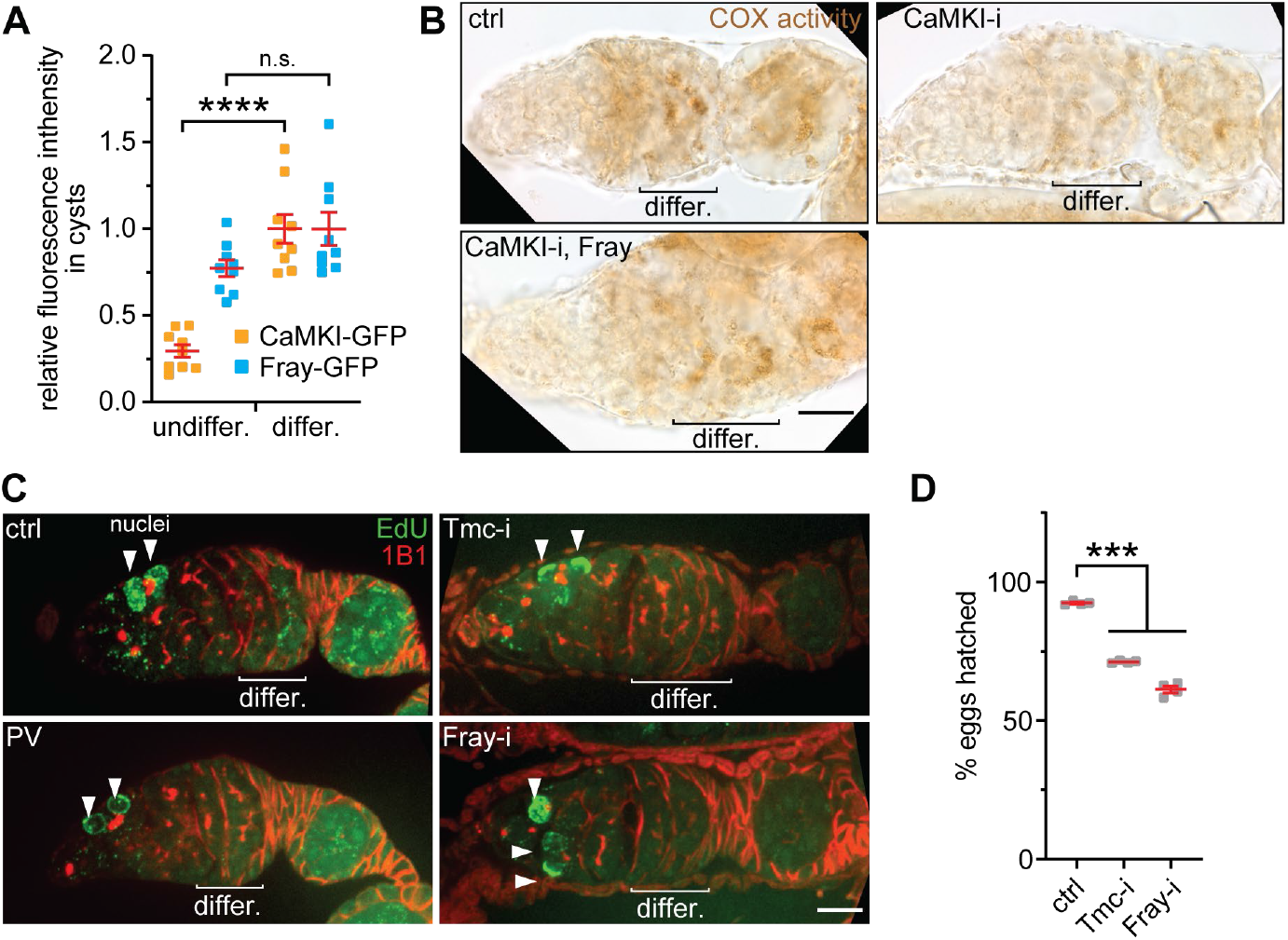
Fray is downstream of CaMKI and the mechanotransduction pathway is required for mtDNA replication in the germline. (**A**) Quantification for the expression levels of endogenous CaMKI-GFP and Fray-GFP in cysts. n = 10 germaria for each protein. (**B**) COX activity histochemistry in germaria. (**C**) mtDNA replication in germaria visualized with EdU incorporation. (**D**) Hatching rate of eggs laid by female flies with indicated germline RNAi. n > 180 eggs laid by 4 females for each genotype. \*\*\**P* < 0.001, \*\*\*\**P* < 0.0001, and n.s., not significant [Mann-Witney tests for (**A**); Kruskal-Wallis tests for (**D**)]. Scale bars, 10 µm.

## Supplementary Tables

**Table S1.**
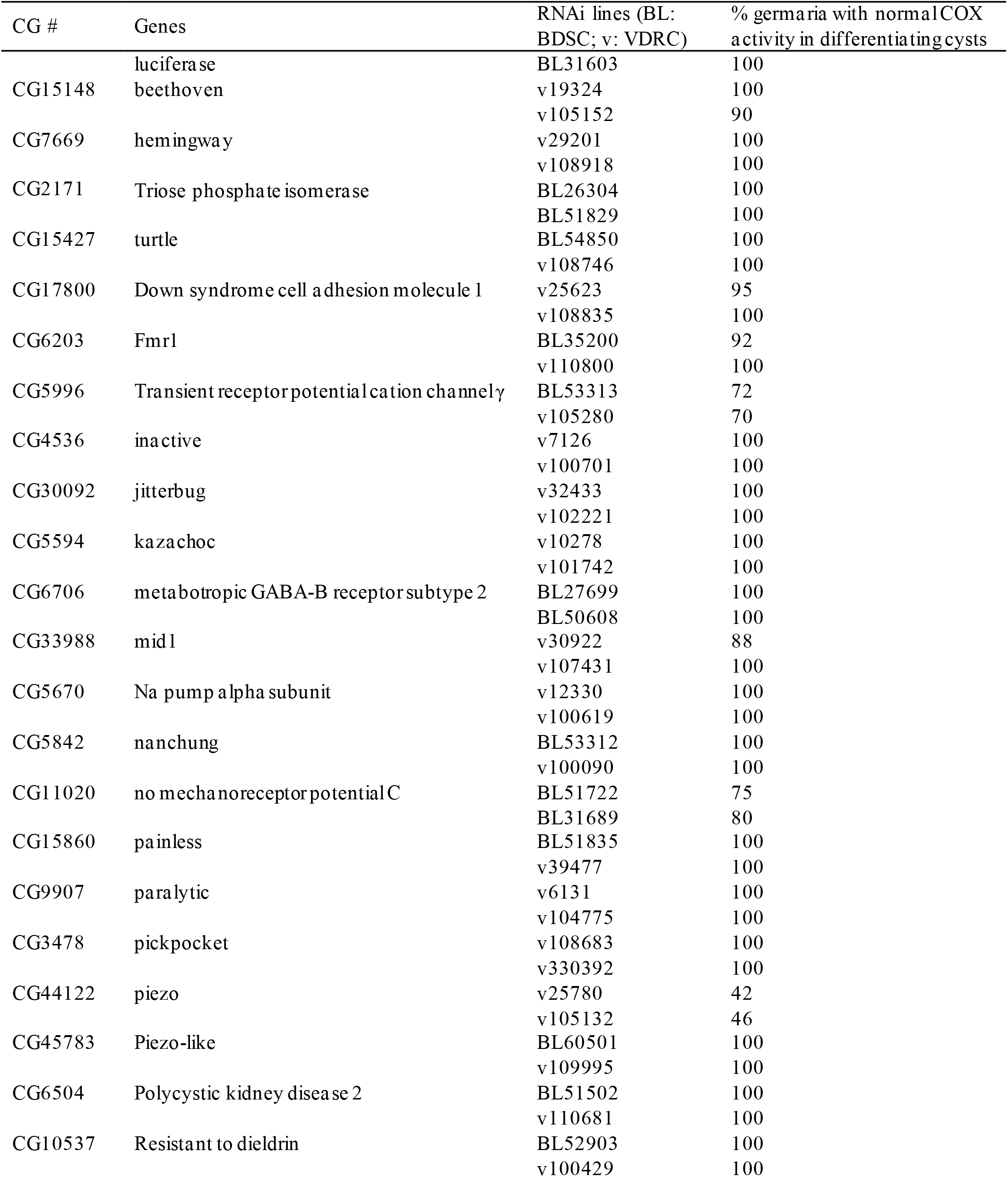
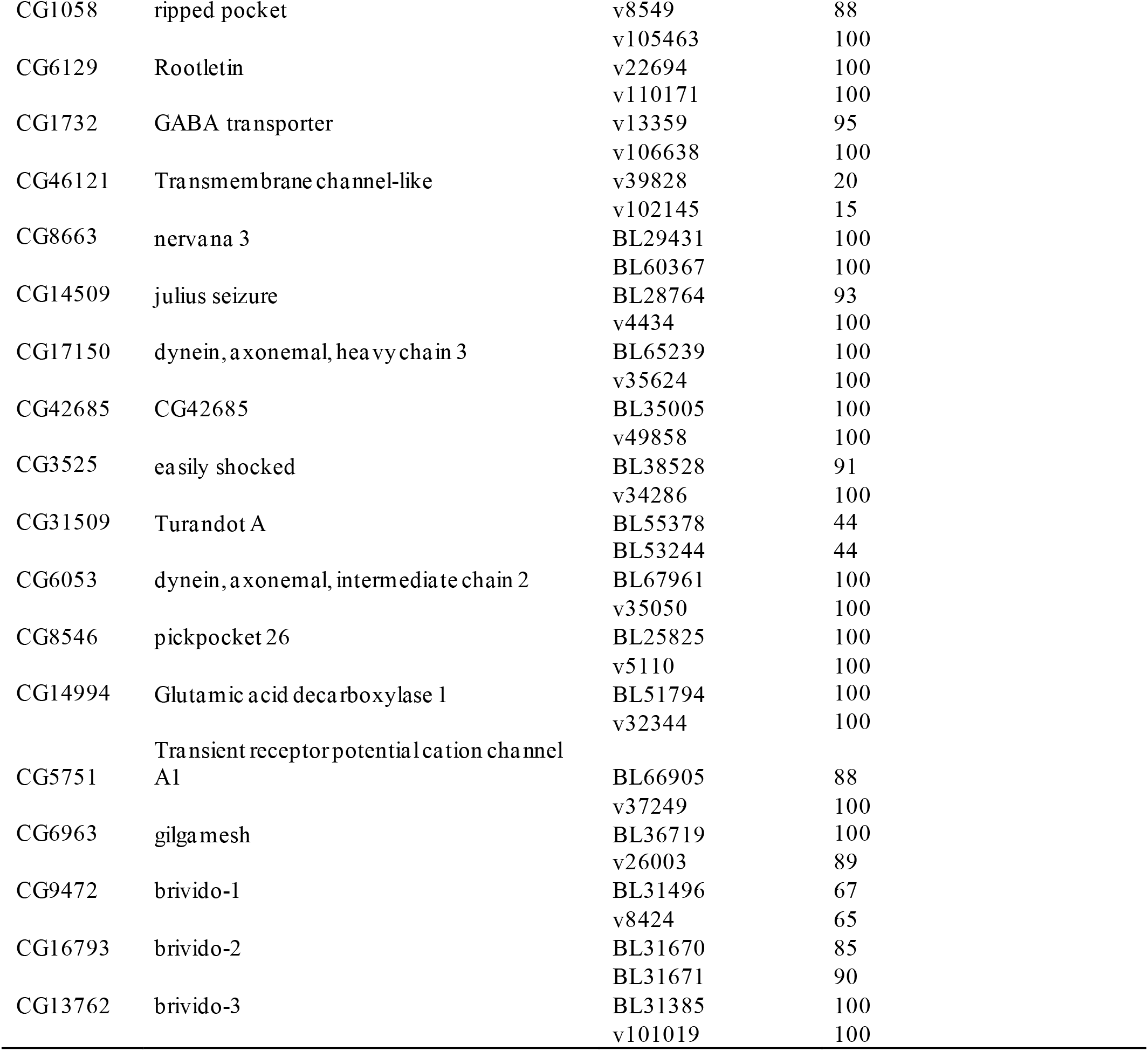
List of RNAi lines for the initial candidate screen used and the resulted COX activity.

**Table S2.**
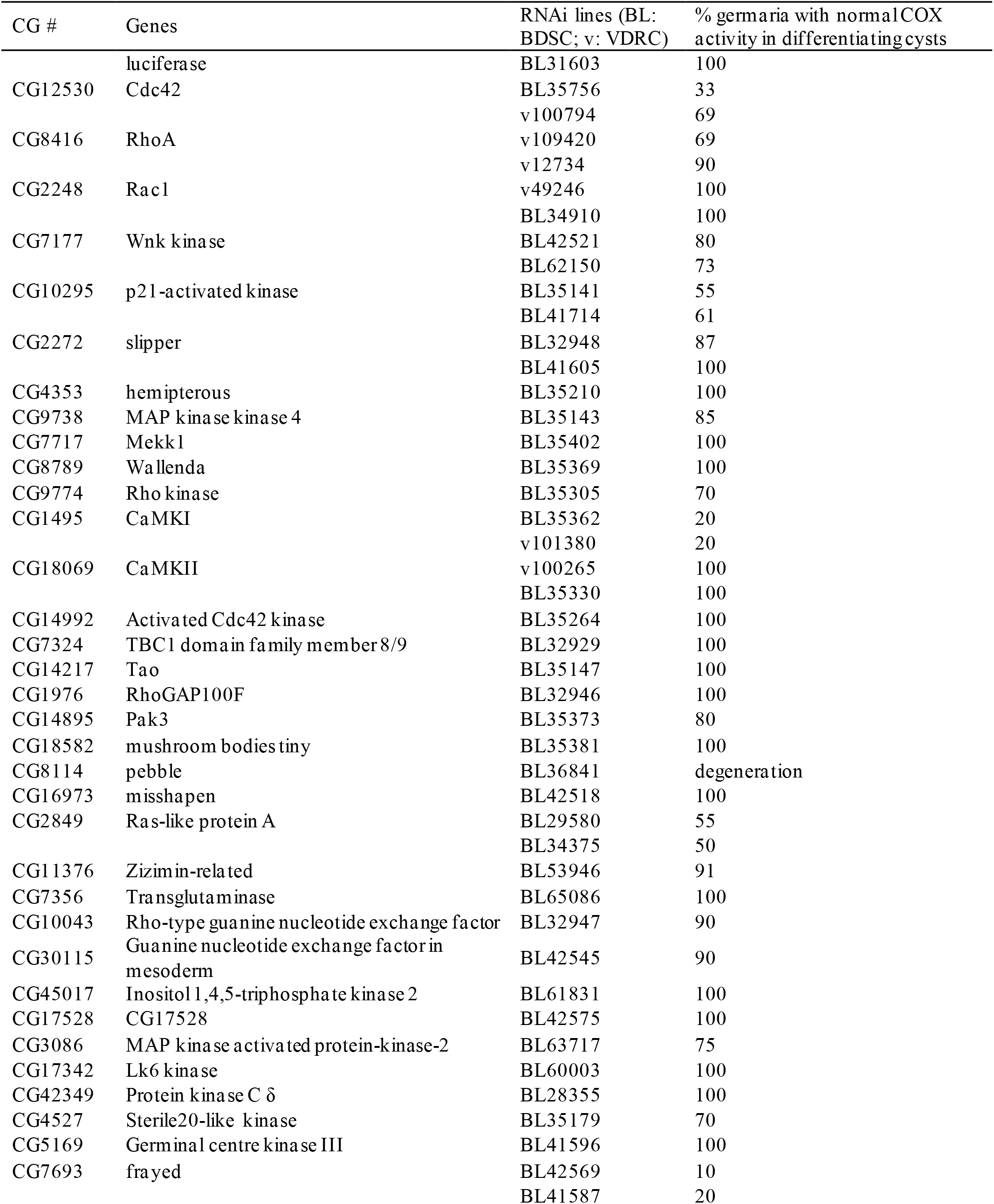
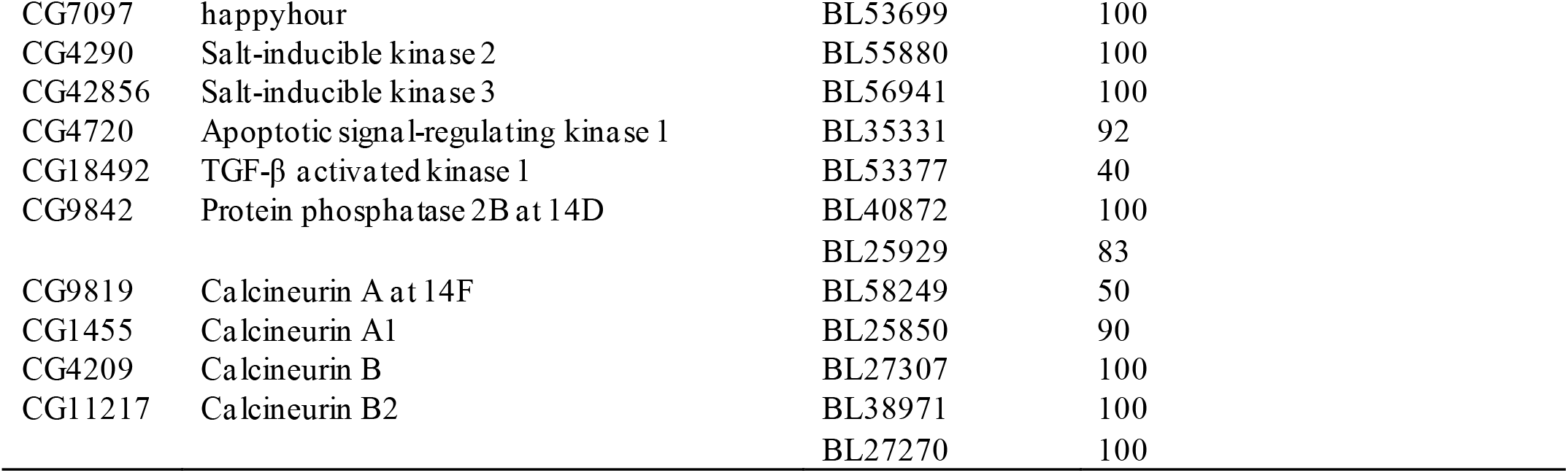
List of RNAi lines for Ca^2+^ signaling and JNK regulators, and the resulted COX activity.

**Table S3.**
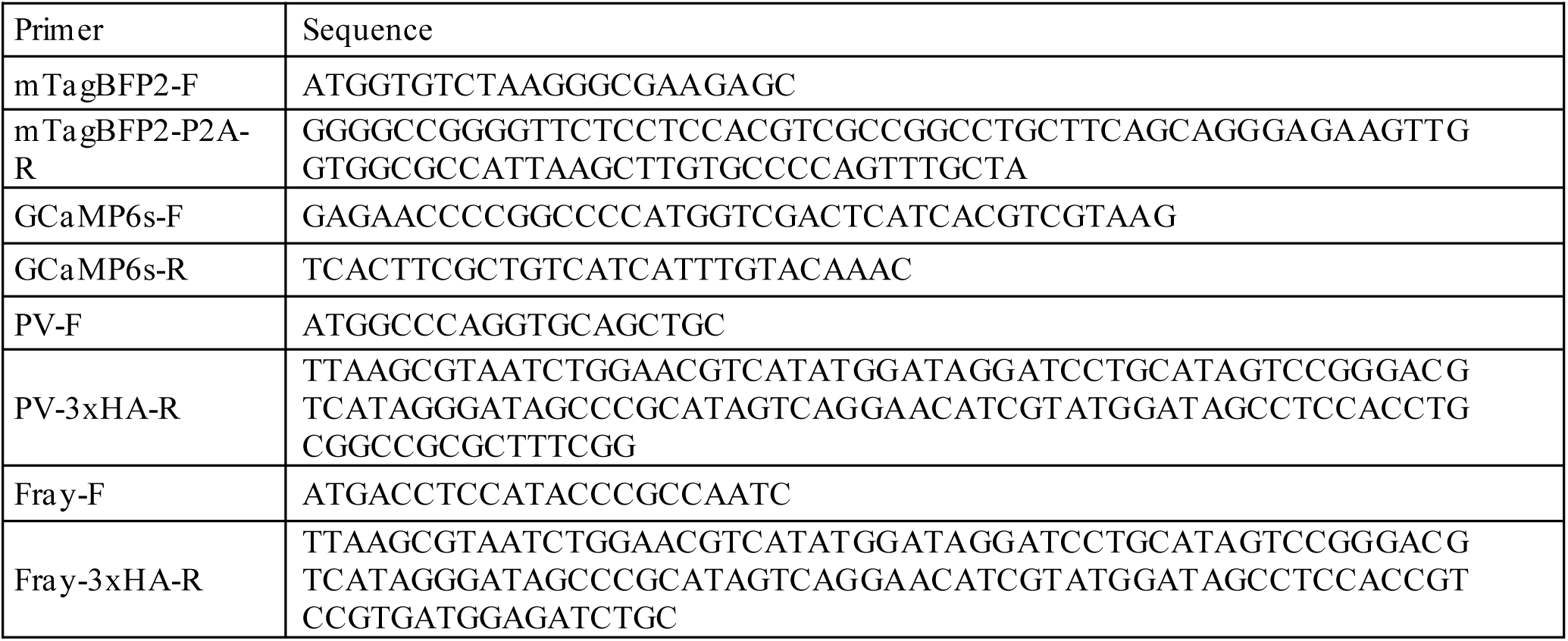
Primers for UASz cloning.

**Table S4.**
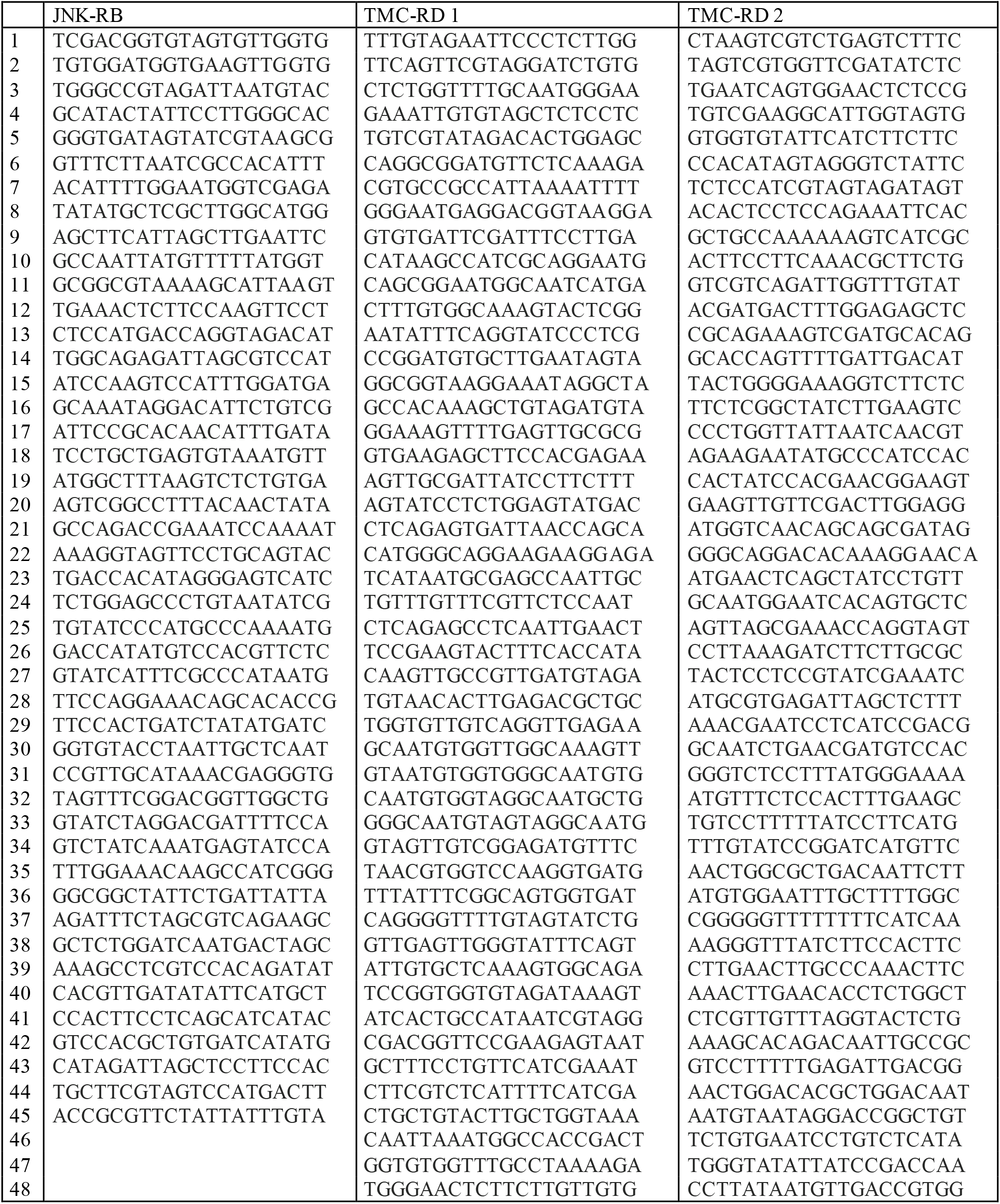
Fluorescently labeled short DNA probes for smFISH.

## Notes

### Competing Interest Statement

The authors have declared no competing interest.

